# Lipid transfer protein ORP3 mediates lysosomal repair via LC3B and ubiquitin-TAK1-p38 signaling

**DOI:** 10.64898/2026.06.09.731146

**Authors:** Christopher J. Bott, Martyna O. Iwaniek, James E. Casanova

## Abstract

Lysosomal membrane damage triggers a multi-stage repair response essential for cellular homeostasis. Here we identify the oxysterol-binding protein-related protein ORP3 as a critical mediator of late-stage lysosomal membrane repair. Following lysosomal damage induced by L-leucine-leucine methyl ester (LLOME) or cationic amphiphilic drugs (CADs), ORP3 is phosphorylated and recruited to ER–lysophagosome contact sites via a signaling cascade initiated by lysosomal membrane ubiquitination, TAK1, p38 MAPK, and, to a lesser extent, IKK. p38-dependent phosphorylation promotes direct interaction between ORP3 and LC3B, which together with PI(4,5)P₂ binding, is required for autophagic lysosome recruitment. ORP3 depletion impairs late-stage lysosomal recovery, elevates lysosomal lipid peroxidation, and reduces cell survival. A lipid transfer-deficient ORP3 mutant fails to restore lysosome function despite normal recruitment, indicating that ER-to-lysophagosome transfer of phosphatidylcholine by ORP3 is functionally required. ORP3 activity is subsequently terminated by VCP/p97-mediated deubiquitination of lysosomes. These findings define ORP3 as a MAPK regulated lipid transfer protein during the late autophagic phase of the endolysosomal damage response.

**Summary:** Lysosomal membrane damage triggers ubiquitination that activates a TAK1-p38 signaling cascade, phosphorylating the lipid transfer protein ORP3 and recruiting it to damaged lysosomes via LC3B interaction. ORP3-mediated phosphatidylcholine transfer from the ER is essential for late-stage lysosomal repair and cell survival.

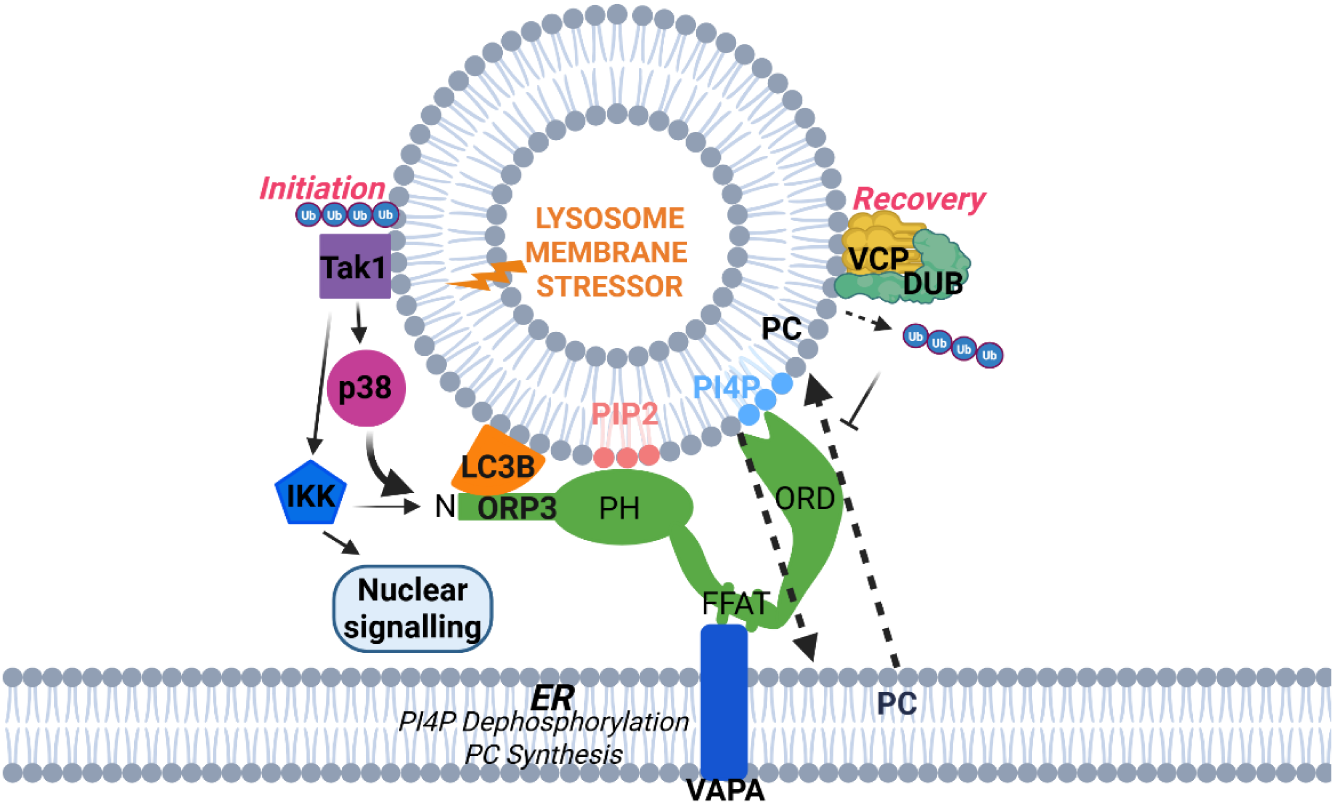

## Introduction

Lysosomes are the primary degradative organelle in cells and are critical for maintaining cellular metabolism, growth, and stress responses. Lysosomal dysfunction is implicated in a broad range of human diseases, including lysosomal storage disorders, pathogen infection, neurodegenerative disease, inflammatory diseases, and aging. Lysosomal membrane integrity can be compromised by the accumulation of undegraded proteins or lipids, reactive oxygen species (ROS), amyloids, bacterial pathogens, and lysosomotropic drugs or toxins (Xun and Tan, 2025; Meyer and Kravic, 2024). Sterile lysosomal membrane damage is commonly modeled using the dipeptide L-leucine-leucine methyl ester (LLOME), which accumulates in acidic compartments via its lysosomotropic properties, where cathepsin C catalyzes its polymerization into membrane-disrupting amyloid-like polymers that cause lysosomal membrane permeabilization (LMP) (Thiele and Lipsky, 1990; Radulovic et al., 2018; Li et al., 2026 *preprint*; Elias et al., 2025 *preprint*). Damaged lysosomes act as signaling organelles that trigger multiple cellular recovery pathways, collectively referred to as the endolysosomal damage response (ELDR) (Meyer and Kravic, 2024). Failure to restore functional lysosomes can trigger inflammation and cell death, and the accumulation of damaged lysosomes is a prominent feature of aging (Tan and Finkel, 2023).

To counteract lysosomal damage, cells employ several repair machineries, each responding to distinct phases of membrane damage. The early, rapid repair stage, occurring within the first 30 minutes, involves recruitment of ESCRTs (Radulovic et al., 2018; Skowyra et al., 2018), bridge-like lipid transfer proteins (BLTPs) (Wang et al., 2025b; Hanna et al., 2025 *preprint*), and the recently described PITT pathway, in which recruitment of the PI4-kinase PI4K2A promotes engagement of lipid transfer proteins including Oxysterol Binding Protein (OSBP) and OSBP-related proteins (ORPs) that function at ER/lysosome contact sites (Tan and Finkel, 2023).

Lysosomal ubiquitination begins at these earliest stages (Gahlot et al., 2024; Eapen et al., 2021; Endo et al., 2025a) and contributes to ESCRT recruitment, membrane repair, and microlysophagy (Zhu et al., 2017; Korbei, 2022)). When rapid repair fails, ubiquitination of lysosomal membrane components escalates, ultimately driving clearance of damaged lysosomes by selective macroautophagy, a process known as lysophagy (Papadopoulos et al., 2017; Reinders et al., 2025; Skowyra et al., 2018; Jia et al., 2020; Li et al., 2026 *preprint*)

Both signaling-associated K63-linked and proteasome-directed K48-linked ubiquitination can occur on membrane proteins of LLOME-damaged lysosomes (Koerver et al., 2019). Beyond recruiting canonical lysophagy adaptors such as p62 and TAX1BP1 (Gallagher et al., 2024), K63 ubiquitination also serves as a signal propagator without substrate degradation (Kiss et al., 2025). On damaged lysosomes, K63 ubiquitin directly recruits and activates the pro-survival MAP3 kinase TAK1, which in turn triggers downstream MAPK and IKK signaling and inflammatory NF-κB activation (Endo et al., 2025a; Celik et al., 2025; Zein et al., 2025; Kanayama et al., 2004). The K63 ubiquitin E3 ligases ITCH and TRIM16 localize to damaged lysosomes in response to distinct damage signals: ITCH is recruited via the lipid-packing defect sensor SPG20, while TRIM16 is recruited by Galectin-3 (Gal3) recognition of exposed luminal glycans on more extensively damaged lysosomes (Gahlot et al., 2024; Chauhan et al., 2016).

Ubiquitination is initiated within minutes of lysosomal damage, increases progressively, and peaks approximately 3 hours after LLOME treatment (Papadopoulos et al., 2017), at which point it begins to be cleared by the ubiquitin-directed AAA-ATPase VCP/p97 in complexes with deubiquitinases (DUBs) such as ATXN3 (Reinders et al., 2025) and USP9X (Jia et al., 2020).

Like OSBP, the ORP family member OSBPL3 (ORP3) functions at ER contact sites through an interaction between its FFAT domain and the ER resident proteins VAP-A and VAP-B (Lehto et al., 2005). We recently reported that ORP3 is recruited to the plasma membrane (PM) through interaction of its PH domain with PI(4,5)P₂ (PIP_2_). We found that the ORP3 ORD domain, which mediates lipid transfer, extracts PI4P from the PM in exchange for ER-derived phosphatidylcholine (PC) (D’Souza et al., 2020). PC is the most abundant phospholipid in cell membranes, and loss of PC synthesis machinery results in autophagosomal maturation defects in yeast (Andrejeva et al., 2020; Polyansky et al., 2022; Orii et al., 2021). Besides PC abundance, PC/phospholipid acyl chain composition has emerged as a major determinant of susceptibility to neurodegenerative disease, cancer, oxidative metabolic disorders, and kidney and liver disease (Estes et al., 2021; Marien et al., 2016; Saum et al., 2026; Nagarajan et al., 2021; Kalyesubula et al., 2025). More recently, ORP3 has been implicated in colorectal cancer progression in part through modulation of MAPK inhibitor sensitivity (Zhong et al., 2026), the nuclear transfer of HIV during infection (Santos et al., 2023), the development of fatty liver disease (Aibara et al., 2023), and metabolic changes following a high-fat diet (Wang et al., 2025a; Kalyesubula et al., 2025; Saum et al., 2026).

ORP3 belongs to the OSBP subfamily III, containing ORP3, ORP6 and ORP7, defined by shared genomic organization and amino acid homology (Lehto et al., 2004; Lehto and Olkkonen, 2003). Among OSBP-related proteins, only ORP3, ORP6, ORP7, and yeast Osh3 form a monophyletic ancient clade tracing back to the last common ancestor of animals and fungi (Singh et al., 2023). ORP3 has a broad tissue distribution and is distinguished by mRNA enriched in HeLa cells, kidney, lymph nodes, and airway, gastrointestinal, and endocrine tissues. In contrast, its close relative ORP6 is primarily expressed in brain and skeletal muscle (Lehto et al., 2004) but is notable due to its localization to damaged neuronal lysosomes and its participation in amyloid-β processing, autophagic lysosome clearance, spinal cord injury recovery, and protection against demyelination. Notably, ORP6 depletion also profoundly disrupts lipid metabolism in brain tissue, including phospholipid acyl chain composition (Zhang et al., 2026; Kasongo et al., 2025; Chang et al., 2026).

ORP3 has recently been implicated in the cellular response to LLOME in several unbiased phosphoproteomic and proteomic studies (Bhattacharya et al., 2023; Jia et al., 2022; Endo et al., 2025b; Su et al., 2026 *preprint*), though it was not the primary focus of those investigations.

Here, we set out to characterize the role of ORP3 in lysosomal repair and recovery. This work is further motivated by accumulating evidence that PIP₂ generation on damaged lysosomes, which is required for ORP3 target membrane engagement, is critical for completion of the late stages of lysophagy (McGrath et al., 2021; Dai et al., 2019; Rong et al., 2012; Tan et al., 2015; Palamiuc et al., 2020)

## Results

### ORP3 is activated and recruited to autophagic lysosomes after LLOME-induced damage, in a p38 MAPK-dependent manner

To determine whether ORP3 associates with damaged lysosomes, we performed live cell imaging of HeLa cells expressing GFP-ORP3. In untreated cells, GFP-ORP3 was diffusely cytosolic; however, it began to slowly condense into discrete puncta following LLOME treatment, which continued to increase in intensity during the ensuing 3 hours (Fig. 1A, B).

**Figure 1:**
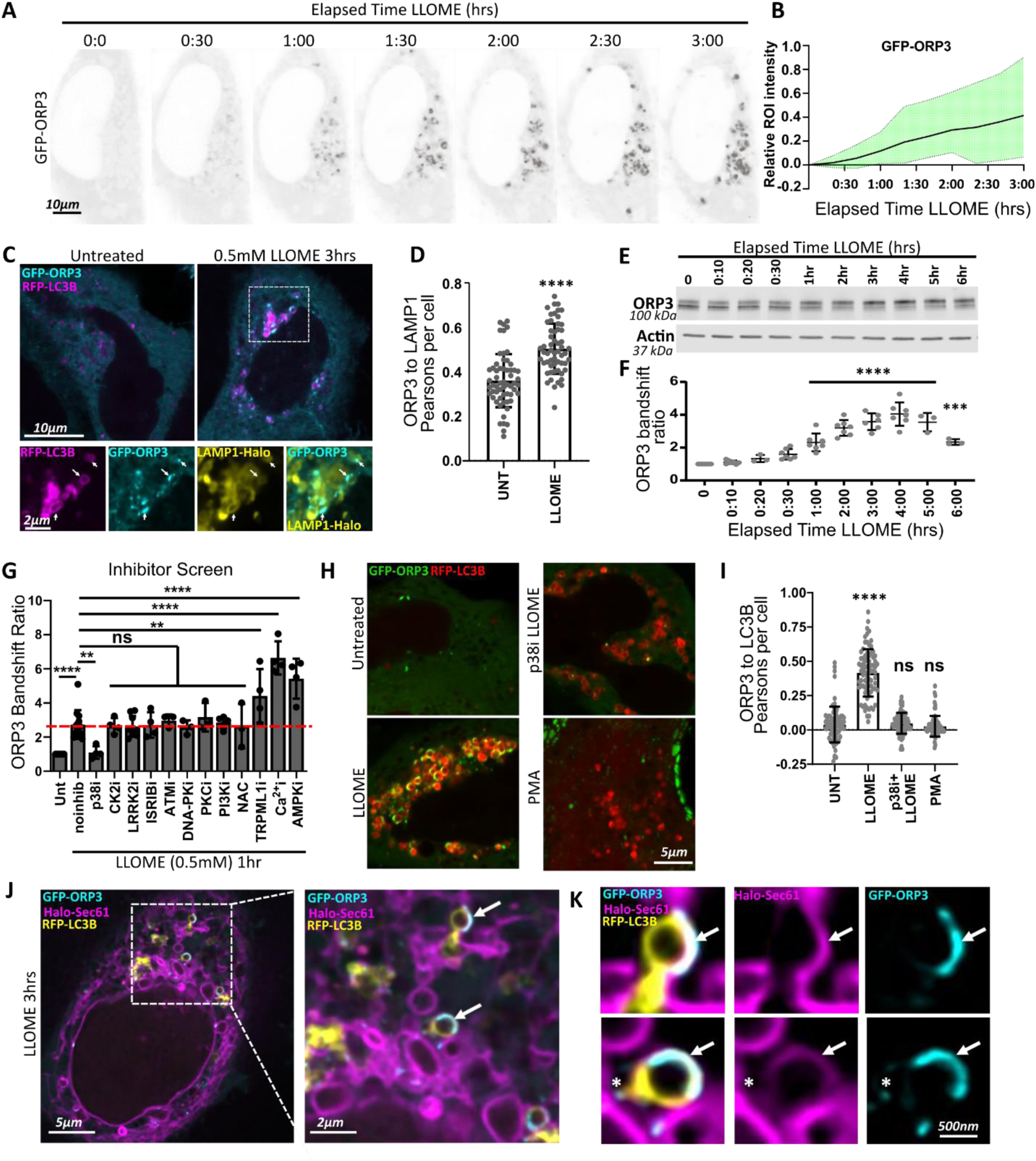
ORP3 translocates autophagic lysosome-ER contact sites in a p38 MAPK-dependent manner at a late stage of lysosomal damage. (**A**) HeLa cells transiently expressing GFP-ORP3 and RFP-LC3B (not shown) were treated with 0.5mM LLOME and imaged live by spinning disc microscopy for 3 h (frames acquired every 10 min). (**B**) Quantification of ORP3 relative intensity over time. n=10 ROIs quantified over 3 experiments. Range indicated by shaded region. (**C**) HeLa cells transiently expressing GFP-ORP3, RFP-LC3B, and Halo-LAMP1 were subjected to LLOME treatment for 3hrs or not, and imaged live via confocal microscopy. Insets from LLOME treated cells. Arrows indicate perinuclear ORP3+ autophagic lysosomes (LAMP1+, LC3B+). (**D**) Quantification of GFP-ORP3 colocalization (Pearson’s coefficient) with immunostained LAMP1 after LLOME 3 h treatment. n=∼60 cells per condition over 3 experiments. Students t-test. **(E)** HeLa cells were treated with the minimal effective dosage of LLOME (0.5mM) and lysed at the indicated times after treatment. Immunoblotting of endogenous ORP3 indicates peak band shift at 3-5 hours. **(F)** Quantification of **E**. n=3 to 7 experiments. One-way ANOVA with Dunnett’s correction tests vs untreated (Time 0). (**G**) Quantification of inhibitor screen of known stress pathways activated downstream of 0.5mM LLOME for 1 hour. Untreated (UNT, vehicle alone)- *Inhibitors*: p38 MAP Kinase (p38i, *Doramapimod)*, Casein Kinase 2 (CK2i, *IQA*), LRRK2 (LRRK2i, *LRRK2-in-1*), Integrated Stress Response (ISRIBi, *Trans-ISRIBi*), ATM Kinase (ATMi, *Ku-55933)*, DNA-PK (DNA-PKi, *NU 7441)*, Protein Kinase C (PKCi, *Go 6983)*, PI3-Kinase (WTMN, *wortmanin)*, Oxidative stress (NAC, *N-acetyl-Cysteine)*, TRPML1/MCOLN (TRPML1i, *ML-SI1)*, Calcium (Ca^2+^I, *BAPTA-AM)*, AMP Kinase (AMPKi, *Dorsomorphin).* Red line indicates the LLOME induced band shift in the absence of inhibitors. N=3 to 12 experiments. One-way ANOVA with Dunnett’s correction tests compared to LLOME with no inhibition. (**H)** Live imaging of HeLa cells transiently expressing GFP-ORP3 and RFP-LC3B and either untreated, treated with LLOME alone, LLOME in the presence of doramapimod (p38i), or with the PKC activator PMA for 3 h. **(I)** Quantification of **H**. n= ∼80 cells over 3 experiments. One-way ANOVA with Dunnet correction tests vs untreated. **(J)** HeLa cells transiently expressing GFP-ORP3, RFP-LC3B, and Halo-Sec61 (ER marker) were treated with LLOME for 3 hours prior to imaging with a Nikon SoRa Super-resolution spinning disc microscope. Arrows in inset indicate ORP3+ autophagic structures in close proximity to Sec61+ ER. (**K**) Higher magnification views of ORP3+ structures indicated by arrows in **J**. Asterisk indicates ORP3+ tubular extension of LC3B+ membranes.

Lysosome disruption by LLOME triggers an autophagic response that can be visualized by tracking recruitment of the autophagic adaptor LC3B to autophagosomal and autolysosomal membranes. In untreated cells, both GFP-ORP3 and RFP-LC3B were diffusely cytosolic (Fig. 1C). After 3 hours of LLOME treatment, both proteins condensed onto LAMP1-immunostained structures resembling swollen autolysosomes or lysosome-containing autophagosomes (lysophagosomes), resulting in significant colocalization between ORP3 and LAMP1 (Fig. 1C, D). Live imaging of Halo-tagged LAMP1 and internalized/chased fluorescent dextran showed comparable colocalization with GFP-ORP3 after LLOME treatment, whereas the late endosomal marker Halo-Rab7 did not (Fig. S1A, B). ORP3 positive organelles typically had a perinuclear, clustered distribution.

Previous work by us and others has shown that ORP3 is phosphorylated by protein kinase C (PKC), promoting its recruitment to ER/plasma membrane (ER-PM) contact sites (D’Souza et al., 2020; Weber-Boyvat et al., 2015; Gulyás et al., 2020). ORP3 displays a reproducible ∼10 kDa band shift on SDS-PAGE gels in response to certain activating stimuli (D’Souza et al., 2020); (Weber-Boyvat et al., 2015; Lehto et al., 2008; Santos et al., 2023; Vertueux et al., 2025 *preprint*), which can be quantified as a ratio of the upper to lower band. Accordingly, “activation” and “band shift” are used interchangeably here. Following LLOME treatment, we observed a significant band shift of endogenous ORP3 in HeLa cells following LLOME treatment, occurring in a dose-and time-dependent manner compared to untreated controls or treatment with L-leucine methyl ester (LOME), a structural analog that does not induce lysosomal damage and serves as a negative control (Fig. 1E, F; Fig. S1C, D). This band shift reflects phosphorylation, as it was reversed by lambda phosphatase treatment of cell lysates regardless of whether it was induced by the Ca²⁺/PKC agonist thapsigargin or by LLOME (Fig. S1E). We found that the minimum effective LLOME dose in HeLa cells at 1 hour was 0.5 mM (Fig. 1F. Fig. S1D), which was used for all subsequent HeLa experiments. The dose response curve was comparable whether LLOME was washed out after 1 hour or maintained throughout the time course, with peak ORP3 activation occurring 3–4 hours after initiating treatment (Fig. 1E, F; Fig. S1F, G). HEK293T cells overexpressing GFP-ORP3 exhibited a similar phosphorylation time course but required 1 mM LLOME to achieve a comparable response (Fig. S1H, I).

To identify the signaling pathway responsible for LLOME-induced ORP3 phosphorylation, we screened a panel of inhibitors targeting known or suspected LLOME-and stress-induced signaling pathways. Cells were pretreated with each inhibitor for 1 hour prior to 1 hour of LLOME incubation. Of the 12 inhibitors tested, only the p38 MAPK inhibitor doramapimod (DMPD) blocked LLOME-induced ORP3 phosphorylation (p38i in Fig. 1G). Two additional p38 inhibitors selective for p38α/β isoforms also effectively blocked ORP3 activation with similar potency (Fig. S1J, K). p38 activation in response to LLOME, as well as activation of other MAPKs, has been previously reported as critical for lysosomal recovery (Gallagher et al., 2024); (Endo et al., 2025a). However, inhibitors of the related MAPKs JNK and ERK1/2 had no effect on LLOME-induced ORP3 activation (Fig. S1L, M). Notably, an inhibitor of LRRK2, which has been implicated in multiple aspects of lysosome repair (Bentley-DeSousa et al., 2025; Bonet-Ponce et al., 2024), did not affect ORP3 phosphorylation (Fig. 1G). In contrast, inhibitors of rapid lysosomal repair mechanisms (Ca²⁺, TRPML1, AMPK), which can exacerbate LLOME-induced damage (Li et al., 2026 *preprint*; Radulovic et al., 2018; Jia et al., 2020), caused increased ORP3 activation.

Importantly, LLOME-induced colocalization between ORP3 and LC3B was completely abolished by DMPD, confirming that p38-dependent phosphorylation of ORP3 drives its recruitment to autophagic lysosomes (Fig. 1H, I). Interestingly, the PKC agonist PMA induced ORP3 translocation to the plasma membrane as previously reported, without triggering colocalization with LC3B (Fig. 1H, I). p38 inhibitors blocked LLOME-induced, but not PMA-induced, ORP3 phosphorylation, while PKC inhibitors blocked PMA-induced phosphorylation but not that triggered by LLOME (Fig. S1N, O). Together, these data indicate that p38 and PKC recruit ORP3 to distinct cellular locations depending on the source of stimulation.

Like other ORPs, ORP3 acts at ER/membrane contact sites (MCS). Super-resolution live imaging (Nikon SoRa) of cells co-expressing GFP-ORP3, RFP-LC3B, and the ER marker Halo-Sec61 confirmed that GFP-ORP3 localizes to the ER on the limiting membrane of swollen autophagic lysosomes (Fig. 1J, K, arrows), often as discrete foci on LC3B-positive tubular extensions (Fig. 1K, white asterisk). Consistent with this, LLOME treatment induced an approximately two-fold increase in co-immunoprecipitation between endogenous ORP3 and GFP-VAP-A, with preferential enrichment of the upper, phosphorylated ORP3 band (Fig. S1P, Q, red arrow). This increase was comparable in magnitude to that induced by the positive control thapsigargin. Importantly, pretreatment with DMPD reduced co-precipitation to basal levels, indicating that the enhanced ORP3/VAP-A interaction is p38-dependent and induced by LLOME.

Finally, we previously reported that ORP3 is recruited to target membranes through interaction of its PH domain with PIP₂, a phosphoinositide typically most abundant at the plasma membrane. Using the PIP₂ biosensor GFP-PLCδ-PH, we confirmed recent evidence that PIP₂ is generated on lysosomes in response to damage (Reinders et al., 2025; Bhattacharya et al., 2023; Posor et al., 2022; Bonet-Ponce et al., 2024) (Fig. S1R).

### ORP3 recruitment is distinct from and sequential to the PITT pathway

The initial PI4P driven phase of ER-contact-mediated lysosomal repair, occurring within the first 30–40 minutes of damage, has been termed the phosphoinositide-initiated membrane tethering and lipid transfer (PITT) pathway (Tan and Finkel, 2022). Lysosomes that cannot be repaired through these mechanisms, likely representing more extensive or stochastic damage, are largely cleared via autophagic engulfment, a process known as lysophagy (Hasegawa et al., 2015; Hoyer et al., 2022; Gahlot et al., 2024).

To determine whether ORP3 is integrated with or distinct from the PITT pathway, we compared the timing of ORP3 recruitment to that of the PITT component ORP11. Consistent with previous studies, we observed significant recruitment of ORP11 to LAMP1-positive lysosomes within 25 minutes of LLOME treatment (Fig.2A, C). In contrast, ORP3 was not detectable on lysosomes at 25 minutes but was robustly present at 3 hours (Fig.2A, B), indicating that ORP3 arrives later than PI4P-dependent PITT pathway components, a timing more consistent with PIP_2_ associated autophagic recovery (Reinders et al., 2025). Unlike ORP3, ORP11 recruitment to lysosomes was insensitive to p38 inhibition (Fig. 2A-C). Similarly, PIKfyve inhibition via vacuolin-1 (Vac1), a specific inducer of the PITT pathway that causes lysosome swelling (Huynh and Andrews, 2005; Yang et al., 2025), failed to activate either p38 or ORP3 (Fig. 6A, B).

**Figure 2:**
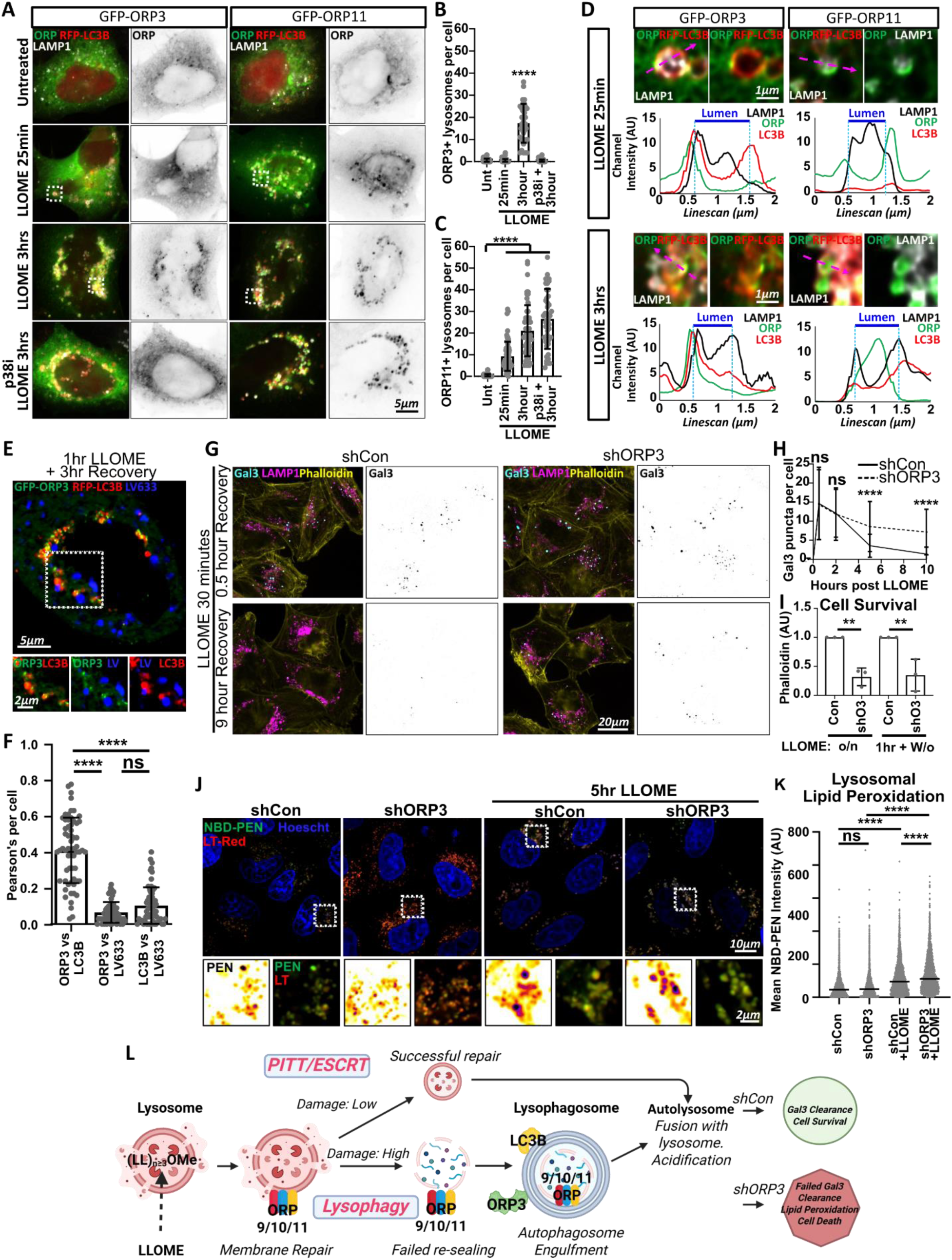
ORP3 recruitment is distinct from and sequential to the PITT lysosomal repair pathway, localizes to non-acidified autolysosomes, and is required for recovery and survival from LLOME (**A**) HeLa cells transiently expressing GFP-ORP3 or GFP-ORP11, as well as RFP-LC3B were treated with LLOME and/or doramapimod (p38i) for the indicated times, fixed, and immunostained for LAMP1, and imaged using conventional confocal microscopy. (**B, C**) Quantification of ORP3+ (**B**) or ORP11+ (**C**) lysosomes at the indicated time points. n= ∼50 cells per condition over 3 experiments. Kruskal-Wallis One-way ANOVA with Dunn’s Multiple Comparisons tests compared to untreated. **(D)** Super-resolution (Nikon NSPARC) imaging of the same cells shown in **A**. Insets of boxed areas in **A** at 25 minutes and 3 hours of LLOME treatment. Line scans are indicated by magenta dotted line. Organelle lumens are indicated by blue and dotted lines in the line scan. **(E)** HeLa cells transiently expressing GFP-ORP3 and RFP-LC3B were treated with LLOME for 1h. The media was changed, and allowed to recover for 3 h prior to imaging. LysoView633 was added 30 minutes prior to imaging. (**F**) Quantification of **E**. Pearson’s colocalization per cell, n= 61 over 3 experiments. One-way ANOVA matched Friedman test with Dunn’s correction as indicated. (**G**) Mock-depleted (shCon) or ORP3-depleted cells (shORP3) were treated with 0.5mM LLOME for 30 min. The media was changed, and allowed to recover for 30 min or 9h, fixed, then immunostained for galectin-3 (Gal3), LAMP1 and actin. (**H**) Quantification of Gal3 puncta at the indicated times after LLOME washout in **G**. n= ∼180 cells per condition over 3 experiments. One-way ANOVA with Sidak’s correction for shCon vs shORP3 at the indicated timepoints. (**I**) Overnight survival of ORP3-depleted HeLa cells (shORP3) vs non-targeting controls (shCon) when treated with 0.5mM LLOME either overnight, or for 1 hour, then washed out. Cells were washed, fixed, and stained with Alexa488-phalloidin. A488 fluorescence relative to untreated controls was measured via plate reader as a readout of cell viability. n=3 experiments per condition. One way ANOVA with Sidak’s correction **(J)** shCon vs shORP3 cells treated with LLOME for 5 h. Live cell imaging of Lysotracker-Red was used to detect lysosomes, while NBD-PEN intensity was used to quantify lipid peroxidation. **(K)** Quantification of **J**. Median NBD-PEN fluorescence per lysotracker+ lysosome. ∼4000-6000 lysosomes per condition over 2 experiments, Kruskal-Wallace One-way ANOVA test. **(L)** Proposed model for early ORP9/10/11 recruitment to lysosomes vs ORP3 recruitment to later autolysosomes, in relation to topology.

We next used super-resolution imaging (Nikon NSPARC) to examine the distributions of ORP3 and ORP11 in greater detail. After 25 minutes of LLOME treatment, ORP11 reliably localized to the limiting membranes of LC3B-negative lysosomes, consistent with its expected function at ER–lysosome contact sites. Interestingly, utilizing the increased power and signal-to-noise ratio of super-resolution imaging revealed subtle changes in ORP3 localization at early-appearing autophagic lysosomes (LAMP1-positive, LC3B-positive) (Fig. 2D) that were not above threshold for conventional confocal microscopy analysis. LAMP1 appeared as smaller puncta within ORP3-positive, LC3B-positive, open-ended, macro-autophagic engulfing membranes (Fig. S2A, A’, A’’), as previously observed by EM (Li et al., 2026 *preprint*). At 3 hours post-LLOME, when the early repair phase has subsided and macro-lysophagy predominates (Su et al., 2026 *preprint*), ORP3 was clearly visible on the rims of autophagic lysosomes (LAMP1-positive, LC3B-positive), consistent with its function at ER contact sites. Surprisingly, ORP11 at 3 hours was predominantly localized to the interior lumen of autophagic lysosomes, suggesting that it had become uncoupled from the ER (Fig. 2D). We hypothesize that lysosomes not repaired by the PITT pathway undergo autophagic engulfment before ORP11 can be retrieved, resulting in this luminal localization (model in Fig. 2L). ORP3-positive structures were largely non-acidified at 3 hours post-LLOME, based on the absence of colocalization with LysoView633 (Fig. 2E, F). This lack of acidification is consistent with a lysophagosome or autophagic lysosome (autophagic engulfment of LAMP1-positive lysosomes) or with immature autolysosomes as reported by others (Reinders et al., 2025)(Fig. 2L).

### ORP3 is required for lysosome recovery after LLOME treatment

Major disruption of the lysosomal membrane allows cytosolic lectins such as Galectin-3 (Gal3) to access exposed luminal glycans, detectable by immunostaining as puncta co-localizing with LAMP1. Loss of these Gal3 puncta over time is widely used as a measure of recovery from lysosomal damage, initially via rapid repair mechanisms, followed by lysophagy (Aits et al., 2015; Meyer and Kravic, 2024). To determine if ORP3 is required for recovery, we depleted cells of endogenous ORP3 using previously validated shRNA (D’Souza et al., 2020) 3 to 4 days before analysis, to avoid major cell compensation.

As shown in Fig. 2G, H, Gal3 was robustly detected on lysosomes in both control and ORP3-depleted cells immediately after LLOME treatment, and the number of puncta remained similar up to 2 hours post-washout during the early stages of lysosomal repair. The ESCRT-associated protein ALIX localized to lysosomes regardless of ORP3 expression, indicating comparable initial damage levels (Fig. S2B, C) and confirming that ORP3 is not required for localization of early lysosomal repair machineries. However, late-stage lysosomal recovery (>2 hours post-washout) was significantly impaired in ORP3-depleted cells. (Fig. 2F, G). This impairment is consistent with the late time course of ORP3 activation (Fig. 1E, F) and resembles recovery deficits reported when other late-stage lysophagy components are disrupted (Bhattacharya et al., 2023; Gahlot et al., 2024). Consistent with a failure to repair damaged lysosomes, ORP3 depletion also reduced cell survival following overnight LLOME treatment (Fig. 2I).

Recently, loss of ORP6, a neuronal-specific close paralog of ORP3, was shown to increase lipid peroxidation following LLOME treatment in HT22 immortalized mouse neuronal cells (Zhang et al., 2026). We hypothesized that ORP3 loss would produce similar effects in HeLa cells, and therefore employed the lipid radical probe NBD-Pen, which has been established as effective for detecting lysosomal lipid peroxidation (Saimoto et al., 2025). NBD-Pen intensity revealed elevated lysosome-localized lipid peroxidation following LLOME in shControl cells, which was significantly exacerbated in shORP3 cells (Fig. 2J, K).

### ORP3 is associated with ubiquitinated lysosomes and is TAK1 dependent

K63 ubiquitination directly drives recruitment of autophagic adaptor proteins that couple damaged lysosomes to autophagic structures for removal by lysophagy. Recent evidence further indicates that ubiquitinated lysosomes also recruit and activate signaling machinery, including the ubiquitin-dependent kinase TAK1 through direct binding of TAK1 complex subunits, TAB1 and TAB2 to polyubiquitin chains (Canovas and Nebreda, 2021; Kanayama et al., 2004). TAK1 is a MAP3 kinase that activates p38 through canonical MAPK signaling (Endo et al., 2025a). A time course of p38 activation following LLOME treatment revealed a biphasic response: an initial pulse during the first 30 minutes, consistent with early ubiquitination events, followed by a second peak at approximately 3 hours (Fig. S3A, B), when ubiquitination reaches its maximum (Papadopoulos et al., 2017). We therefore hypothesized that p38-dependent ORP3 phosphorylation is initiated through K63 ubiquitin-mediated recruitment and activation of TAK1.

To test this hypothesis, we first confirmed that lysosomal membranes become prominently poly-ubiquitinated in response to LLOME, as detected by the poly-ubiquitin-specific (K48 and K63) monoclonal antibody FK2 (Fujimuro and Yokosawa, 2005; Koerver et al., 2019). As previously reported (Papadopoulos et al., 2017), we detected a significant increase in FK2 staining after 3 hours of LLOME treatment (Fig. 3A, B), coinciding with peak ORP3 activation (Fig. 1E, F).

**Figure 3:**
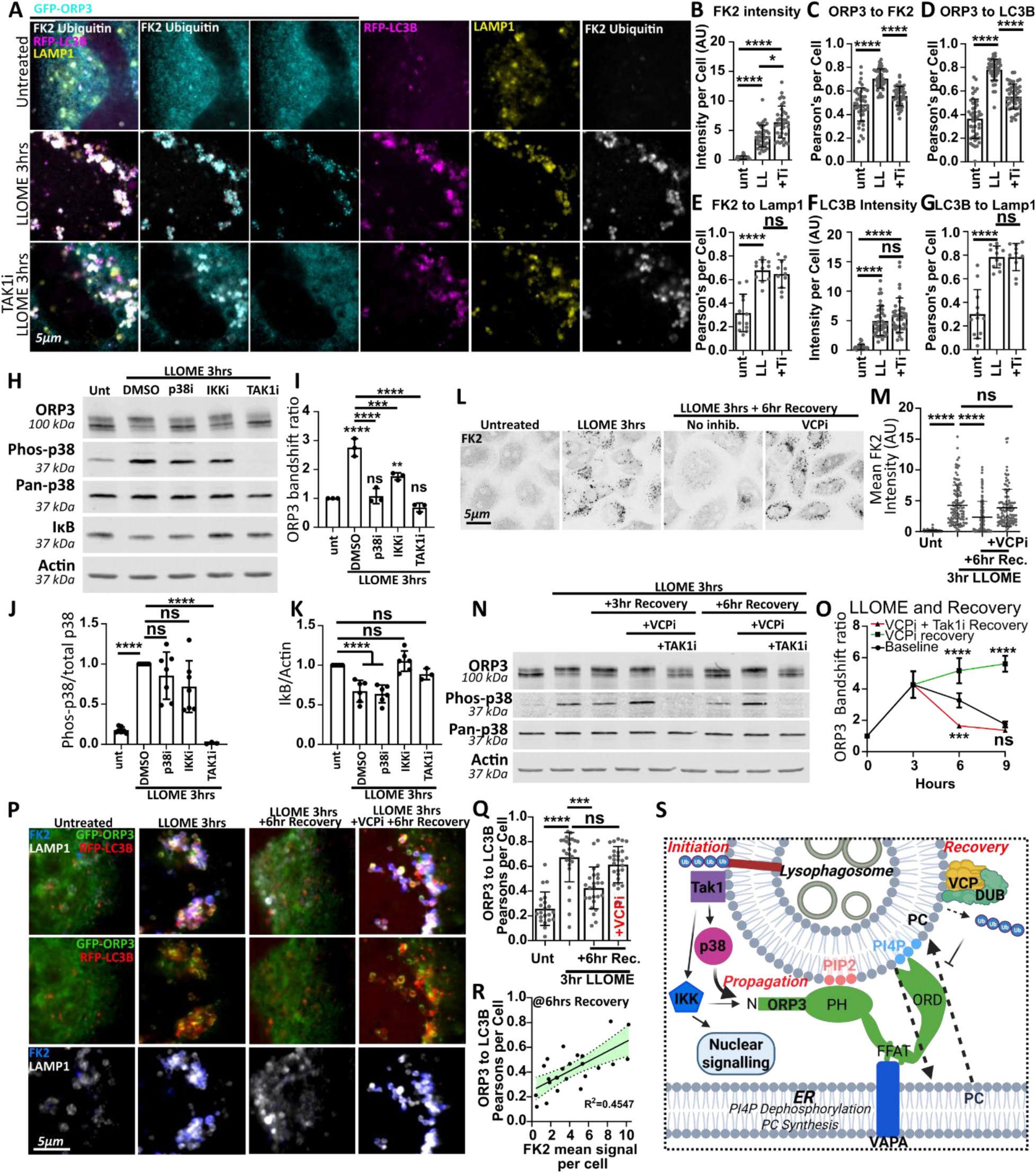
ORP3 activation occurs downstream of ubiquitin-stimulated TAK1 activity, and is deactivated via a deubiquitinase/VCP complex (**A**) HeLa cells transiently expressing GFP-ORP3 and RFP-LC3B were untreated (top row), or treated with LLOME for 3 h in the absence (middle row) or presence of the TAK1 inhibitor (5Z)-7-Oxozeaenol (TAK1i) 30 min prior to LLOME treatment. Cells were then fixed and immunostained for LAMP1 and K48/K63 poly-ubiquitin with monoclonal antibody FK2. (**B-G**) Quantification of **A**. (**B**) n∼40 cells for each condition over 3 experiments, One Way ANOVA with Tukey’s correction. (**C, D**) n∼40 cells for each condition 3 experiments, One-way ANOVA with Sidak’s correction. (**E**) n=12 cells over 1 experiment, One-way ANOVA with Sidak’s correction. (**F**) n∼40 over 3 experiments, One way ANOVA with Tukey’s correction. (**G**) n=12 cells over one experiment, One-way ANOVA with Sidak’s correction. (**H**) HeLa cells treated with LLOME for 3h with the indicated inhibitors were immunoblotted for the indicated targets. (**I**) Quantification of the relative ORP3 bandshift under conditions indicated in **H**. N=3 to 7 experiments One-way ANOVA with Sidak’s correction___(**J**) Densitometric quantification of phospho-p38 (p-T180, p-Y182) relative to total p38 under conditions indicated in **H**. N=3 to 7 experiments, One-way ANOVA with Dunnett’s correction (**K**) Densitometric quantification of cytosolic IкB relative to actin loading control under indicated conditions in **H**. n=3 to 6 experiments One-way ANOVA with Dunnet’s correction **(L)** HeLa cells were left untreated or treated with LLOME for 3h, then media was changed and allowed to recover for 6h in the presence or absence of the VCP inhibitor *NMS-873* (VCPi). Cells were then fixed and stained for K48/K63 poly/mono-ubiquitin with FK2 antibody. (**M**) Quantification of **L**. n=90-130 cells, Kruskal Wallace test with Dunn’s correction. (**N**) HeLa cells were treated with LLOME for 3 hours, fresh media was added and cells allowed to recover in the presence of *NMS-873* (VCPi) or *(5Z)-7-Oxozeaenol* (TAK1i) for the indicated times. (**O**) Quantification of **N**. n=3-5 experiments. One-way ANOVA with Sidak correction for multiple comparisons relative to recovery in the absence of inhibitors. (**P**) HeLa cells transiently expressing GFP-ORP3 and RFP-LC3B were treated or not with LLOME for 3 h, media were changed, and allowed to recover in fresh media for the indicated times in the presence or absence of *NMS-873* (VCPi), fixed, and immunostained for LAMP1 and K48/K63 poly-ubiquitin (FK2). **(Q)** Quantification of Pearson’s colocalization between ORP3 and LC3B as shown in **P**. n∼25 cells, Kruskal-Wallace One-way ANOVA test. (**R**) Linear relationship between the remaining ubiquitin in cells and ORP3/LC3B colocalization per cell as shown in **P**. n∼25 cells, Simple linear regression analysis. (**S**) Model for ubiquitin/TAK1/MAP3K-mediated activation of ORP3, and VCP/DUB (deubiquitinase) clearance of ubiquitin and deactivation of ORP3. Created with BioRender.

ORP3 localized to these FK2-positive autophagic lysosomes and this localization was completely abolished by pretreatment with the TAK1 inhibitor 5Z-7-Oxozeaenol (TAK1i) (Fig. 3A). TAK1 inhibition did not diminish upstream events such as lysosomal ubiquitination (Fig. 3E), LC3B lipidation, or LAMP1/LC3B colocalization, indicating a direct effect on ORP3 activation (Fig. 3 C-G). Immunoblotting confirmed that the TAK1 inhibitor suppressed LLOME-induced activation of both p38 and ORP3 (Fig. 3H–J). MG132, which results in non-specific accumulation of polyubiquitin species including K63 (Heidelberger et al., 2018), also results in TAK1-dependent ORP3 activation similarly to LLOME (Fig. S3C, D).

LLOME-associated TAK1 signaling has also been reported to promote IKKα/β activation, IκB degradation, and subsequent activation of the inflammatory transcription factor NF-κB (Endo 2025). We confirmed that inhibition of either TAK1 or IKKα/β, but not p38, significantly blocked LLOME-induced IκB degradation, placing TAK1 independently upstream of both p38 and IKK (Fig. 3H-K). However, IKKα/β inhibition had only a partial effect on ORP3 phosphorylation (Fig. 3H, I) and its recruitment to autophagic lysosomes (Fig. S3E-G). Inhibitors of TBK1 and ULK1, kinases known to act downstream of TAK1/p38 in the context of autophagy and infection (Ma et al., 2023; Slobodnyuk et al., 2019), did not significantly disrupt ORP3 phosphorylation following LLOME treatment (Fig. S3H, I).

### ORP3 is deactivated by VCP/p97-mediated deubiquitination of lysosomal membranes

Given that lysosomal K63 ubiquitination drives TAK1/p38-dependent ORP3 activity, we also investigated the VCP(p97)/DUB complex, which has been shown to remove K63 poly-ubiquitin from PIP₂-positive, non-acidified lysosomes (Reinders et al., 2025) like those decorated by ORP3. VCP/DUB complex recruitment begins approximately 2 hours after LLOME treatment and persists at later timepoints (Reinders et al., 2025). As shown in Fig. 3L, M, lysosomal ubiquitination decreased significantly after 6 hours of recovery from a 3-hour LLOME treatment. Consistent with previous reports (Reinders et al., 2025), VCP/p97 inhibition blocked ubiquitin removal and recovery. In parallel, phosphorylation of both p38 and ORP3 returned to near-basal levels after 6 hours of recovery in the absence of inhibitors (Fig. 3N, O), and confocal microscopy confirmed a corresponding reduction in ORP3/LC3B colocalization (Fig. 3P, Q).

Although ORP3/LC3B colocalization had not fully returned to baseline at this timepoint, we observed a linear relationship between residual ubiquitin (FK2) signal per cell and ORP3/LC3B colocalization (Fig. 3R), further supporting a link between ubiquitination levels and ORP3 activity. In contrast, when ubiquitin turnover was blocked by VCP inhibition, phosphorylation of p38 and ORP3 (Fig. 3N, O) and ORP3/LC3B colocalization (Fig. 3P, Q) persisted throughout the time course, an effect completely reversed by co-treatment with the TAK1 inhibitor (Fig. 3N, O). Of note, VCP inhibition alone without LLOME had little effect on ORP3 activation (Fig. S3J, K, K). Together, these data indicate that VCP/DUB-mediated deubiquitination of lysosomes terminates TAK1/p38-mediated ORP3 activity (Fig. 3S).

### ORP3 interacts with LC3B in a p38-dependent manner

Several studies have suggested an interaction between ORP3 and ATG8 family proteins, including LC3B (Tu et al., 2022; Bhattacharya et al., 2023). While endogenous ORP3 did not co-precipitate with GFP-LC3B in untreated HeLa cells, the two proteins interacted robustly after either 1.5 or 3 hours of LLOME treatment (Fig. 4A, B). To determine whether this interaction requires p38-dependent ORP3 phosphorylation, we used recombinant purified GST-LC3B to precipitate GFP-ORP3 from lysates of LLOME-treated HEK293T cells in the presence or absence of doramapimod. ORP3 did not interact with unfused GST under any condition (Fig. 4E, F). Consistent with the immunoprecipitation shown in Fig. 4A, ORP3 was barely detectable in GST-LC3B precipitates from untreated cells but was robustly pulled down from LLOME-treated cells, again favoring the upper, phosphorylated, ORP3 band. Importantly, p38 inhibition with doramapimod, which blocked ORP3 phosphorylation, completely abolished co-precipitation with GST-LC3B, indicating that p38-dependent phosphorylation is required for this interaction (Fig. 4E, F). Notably, PMA, which induces robust ORP3 phosphorylation via PKC (Fig. S1N, O), did not promote an ORP3/GST-LC3B interaction (Fig. 4C, D).

**Figure 4:**
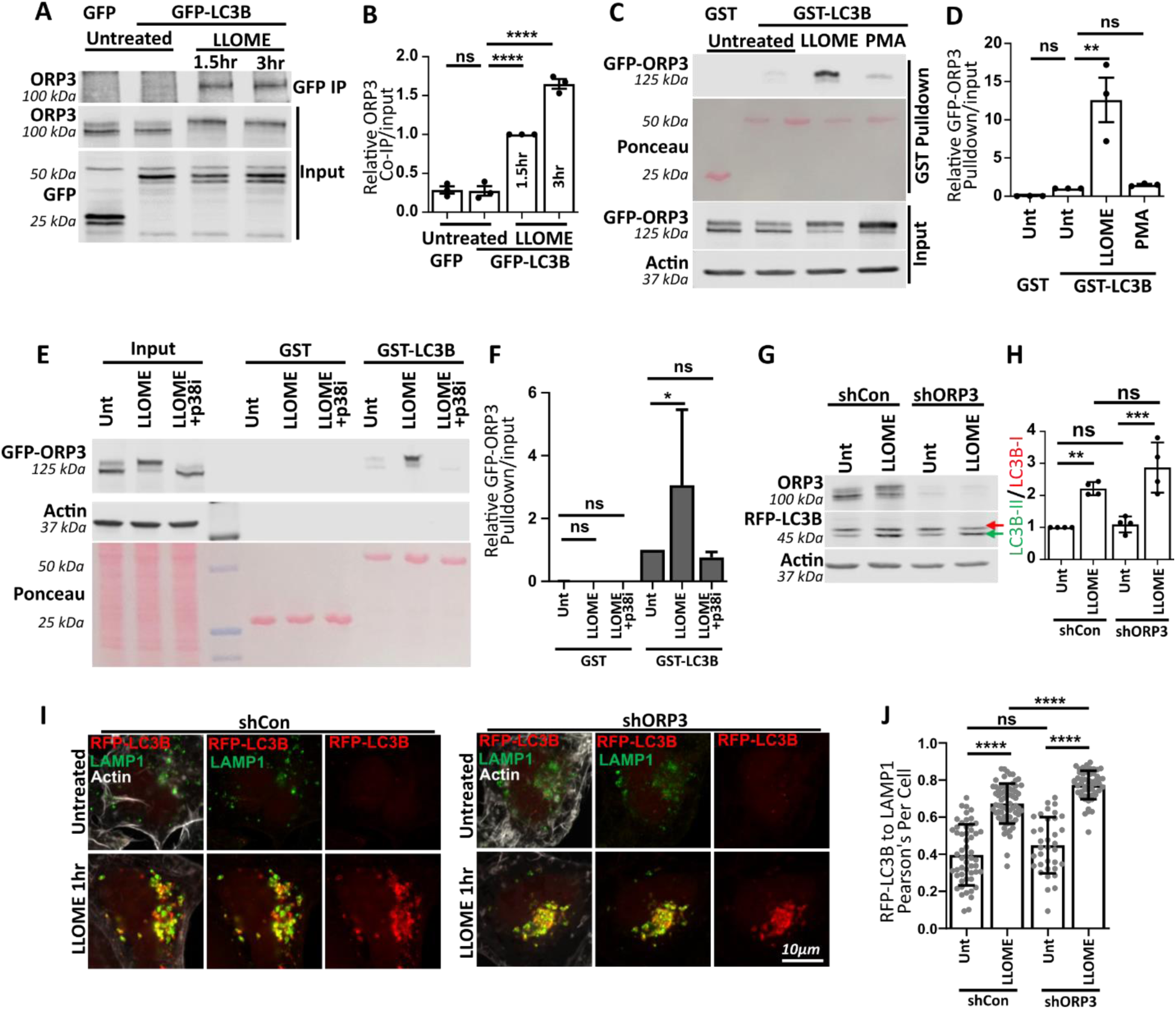
ORP3 interacts with LC3 in a p38-dependent manner. (**A**) HeLa cells transiently expressing GFP or GFP-LC3B were treated for the indicated times, lysed, and immunoprecipitated with an anti-GFP antibody. Co-immunoprecipitation of endogenous ORP3 was assessed via immunoblot. (**B**) Densitometric quantification of **A**, n=3 experiments, One-way ANOVA (**C**) HEK293T cells transiently expressing GFP-ORP3 were treated with LLOME or PMA for 3 h. Cells were then lysed, subjected to pulldown with 4ug of GST alone or 4ug GST-LC3B agarose beads (25 and 50 kDa on Ponceau-stained blot, respectively), and immunoblotted for ORP3 or actin. (**D**) Quantification of **C**, n=3 experiments, One-way ANOVA. (**E**) HEK293T cells transiently expressing GFP-ORP3 were treated with LLOME +/-doramapimod (p38i) for 3h. Cells were lysed and subjected to pulldown with 4ug GST alone or 4ug GST-LC3B agarose beads (stained with Ponceau as above), and immunoblotted for ORP3 and actin. (**F**) Quantification of **E**. n=3-5 experiments, One-way ANOVA (**G**) HeLa cells transiently expressing RFP-LC3B were depleted of ORP3 (shORP3) or not (shCon) and treated with LLOME for 1h, then immunoblotted for endogenous ORP3 and RFP-LC3B. (**H**) Quantification of LC3B lipidation in **G** (densitometry of LC3B-II relative to LC3B-I). n=4 experiments, One-way ANOVA (**I**) HeLa cells transiently expressing RFP-LC3B and depleted of ORP3 were treated with LLOME for 1h, fixed and immunostained for LAMP1. (**J**) Quantification of LC3B/LAMP1 colocalization (Pearsons coefficient) in **I**. n=50-70 cells over 2 experiments, One-way ANOVA.

ORP3 loss did not affect LC3B lipidation in response to LLOME, as conversion to the lipidated LC3-II form proceeded normally in ORP3 depleted cells (Fig. 4G, H). Similarly, LC3B localization to damaged lysosomes occurred at comparable frequencies in control and ORP3-depleted cells (Fig. 4I-J), indicating that ORP3 functions downstream of LC3B, despite the small amounts appearing on pre-autophagic structures (Fig. 2D; Fig. S2A). This suggests that ORP3 functions after autophagosome formation.

### ORP3 interacts with LC3 via an LC3-interacting region (LIR)

Bioinformatic analysis using the iLIR autophagy database (Jacomin et al., 2016) and the Eukaryotic Linear Motif (ELM) (Kumar et al., 2024) resource identified a highly evolutionarily conserved “Type W” LC3-interacting region (LIR) (Rogov et al., 2023) and an adjacent KKRK polybasic motif in the N-terminus of ORP3 (Fig. 5A, B), revealed utilizing COBALT, a constraint based multiple alignment tool (Papadopoulos and Agarwala, 2007). AlphaFold3 modeling of the ORP3–LC3B interaction revealed a canonical ATG8–LIR interaction, in which the hydrophobic residues of the ORP3 LIR insert into hydrophobic pockets 1 and 2 of the LC3B LIR Docking Site (LDS) (Fig. 5A-C) (Abramson et al., 2024; Johansen and Lamark, 2020; Rogov et al., 2023).

**Figure 5:**
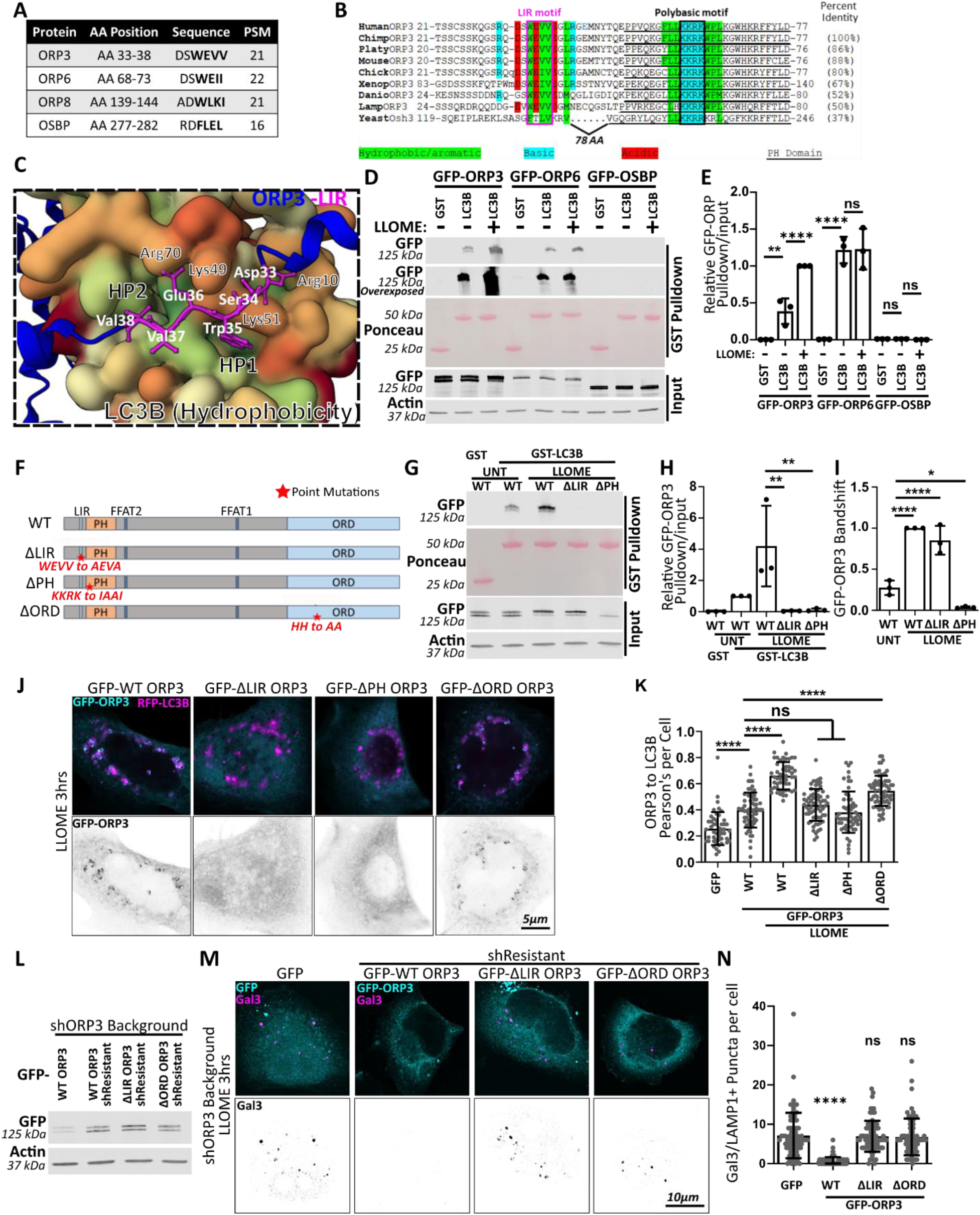
ORP3 directly interacts with LC3 via an LIR domain, and phosphorylation of ORP3 is dependent upon lipid interactions with the PH domain. (**A**) Bioinformatics of putative LC3B Interacting Regions (LIRs) of OSBP/ORP family proteins. (**B**) Constraint-based Multiple Alignment of ORP3 proteins from different organisms. Conserved LIR, polybasic region, and other relevant conserved features are marked as indicated. OSBPL3 sequences from *Homo sapiens* (Human); *Pan troglodytes* (Chimp); *Ornithorhynchus anatinus* (Platypus); *Mus musculus* (Mouse); *Gallus gallus* (Chicken); *Xenopus laevis* (Xenopus), *Danio rerio* (Danio); *Lampetra planeri* (Lamprey); and *Saccharomyces cerevisiae* (Yeast). (**C**) Alphafold3 model of the ORP3-LIR (relevant amino acids in white) and LC3B (Gaussian surface hydrophobicity projection, where more hydrophobic = green) (relevant LC3B amino acids in black). Hydrophobic Pocket 1 and 2 (HP1 and HP2) of LC3B are indicated. (**D**) Lysates of HEK293T cells transiently expressing GFP-ORP3, GFP-ORP6, or GFP-OSBP were incubated with GST-LC3B and immunoblotted for LLOME-dependent interaction. (**E**) Densitometric quantification of precipitated GFP-tagged ORP/OSBP protein relative to input as shown in **D**. n = 3 experiments, One-way ANOVA. **(F)** Schematic diagram of ORP3 domain structure, with description and location of point mutations (red). (**G**) GFP-ORP3 ΔLIR and GFP-ORP3 ΔPH mutants were expressed in HEK293T cells, and assessed for LLOME-dependent binding to LC3B in comparison to WT ORP3. (**H**) Quantification of pulldown efficiency in **G**. n = 3 experiments, One-way ANOVA with Sidak’s correction. (**I**) Quantification of ORP3 mutant phosphorylation (upper band relative to lower band) in **G**. n=3 experiments, One-way ANOVA with Dunnett’s correction. (**J**) HeLa cells transiently expressing GFP-ORP3 WT, ΔLIR, ΔPH, or ΔORD, in addition to RFP-LC3B, were treated with LLOME for 3h, fixed and imaged by confocal microscopy. (**K**) Pearson’s co-localization of ORP3 WT or mutant constructs with RFP-LC3B after LLOME treatment in **J**. n=50-70 cells over 2 experiments, One-way ANOVA with Dunnett’s correction. (**L**) Expression of shRNA-resistant ORP3 rescue constructs in HeLa cells depleted of endogenous ORP3. (**M**) Rescue of ORP3-depleted cells expressing GFP or shRNA-resistant GFP-ORP3 WT, GFP-ORP3 ΔLIR or GFP-ORP3 ΔORD. Cells were treated with LLOME for 30 min, washed and allowed to recover for 8h. Cells were fixed and stained for Galectin-3. (**N**) Quantification of Gal3 puncta per cell in **M**. n=70 to 90 cells over 4 experiments, One-way ANOVA with Dunnett’s correction.

**Figure 6:**
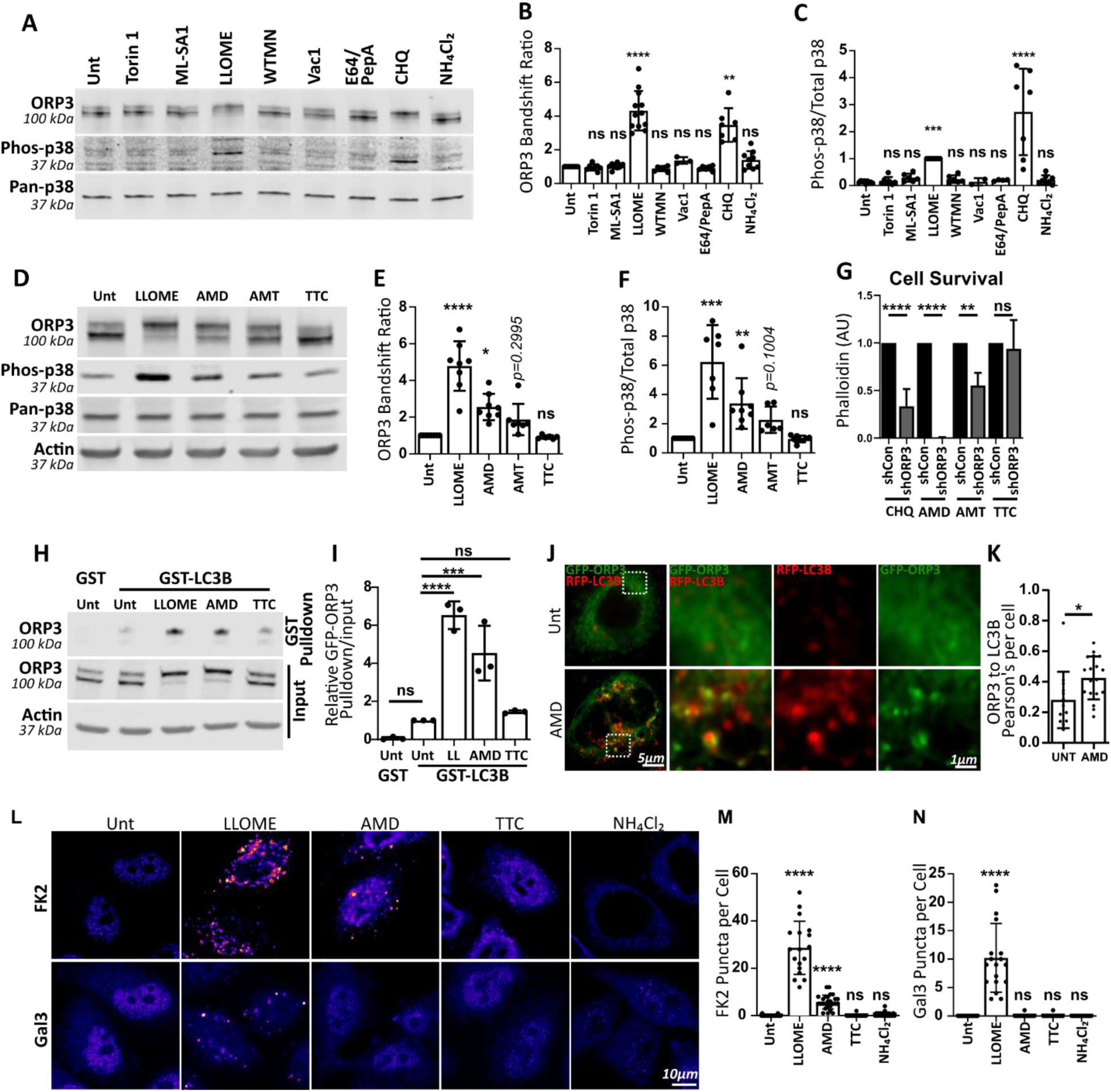
Cationic Amphiphilic Drugs induce activation of p38 and ORP3, and decrease survival in ORP3-depleted cells. (**A**) HeLa cells were treated with the indicated agents for 3h, then immunoblotted to detect endogenous ORP3 band shift and p38 activation. (**B, C**) Quantification of ORP3 and p38 activation by agents shown in **A**. n=3-12 experiments per condition and n=2-11 experiments, respectively, Kruskal-Wallace test. (**D**) HeLa cells were treated with indicated Cationic Amphiphilic Drugs (CADs) for 3 h and immunoblotted to detect endogenous ORP3 band shift and p38 activation. (**E, F**) Quantification of ORP3 and p38 activation by CADs in **D**. n=6-8 experiments per condition, Kruskal-Wallace test. (**G**) Survival of ORP3-depleted HeLa cells (shORP3) vs non-targeting controls (shCon) after treatment with the indicated CADs overnight. Cells were washed, fixed, and stained with Alexa488-phalloidin. A488 fluorescence was measured by plate reader as a readout of cell viability. n=4 to 5 experiments per condition. One way ANOVA with Sidak’s correction. **(H)** HeLa cells expressing GFP-ORP3 were treated with LLOME, amiodarone (AMD), amytryptyline (AMT) or tetracycline (TTC) for 3h, lysed and incubated with GST-LC3B beads, then immunoblotted for ORP3 and actin. **(I)** Quantification of **H**. n=3 experiments per condition, One-way ANOVA relative to untreated GST-LC3B pulldown. **(J)** HeLa cells expressing GFP-ORP3 and RFP-LC3B were treated with AMD for 3 h or left untreated. Boxed areas are enlarged on the right. **(K)** Pearson’s colocalization of ORP3 with LC3B under conditions shown in **J**. n∼15 cells per condition, unpaired t-test. **(L)** HeLa cells were treated with LLOME or the indicated CAD for 3 h, then fixed and stained for polyubiquitin (FK2) or Gal3. **(M, N)** Quantification of FK2 and Gal3 puncta per cell as in **L**. n= 20-30 cells per condition. Kruskal Wallace test compared to untreated.

Interestingly, several other ORP family members also contain predicted LIRs (Fig. 5A). ORP6 also contains a high-confidence LIR. Notably, ORP6 interacted with GST-LC3B constitutively, regardless of LLOME treatment, in contrast to the stimulus-dependent interaction of ORP3 (Fig. 5D, E). Constraint based multiple alignment of ORP6 and ORP3 indicated similar LIR placement immediately adjacent to a conserved KKRK polybasic PIP₂-binding motif (Golebiewska et al., 2006) in their PH domains (Fig. S5A). However, ORP6 contains a 33-residue intrinsically disordered insert immediately upstream of its LIR that is not present in ORP3, which may contribute to their differential interaction with LC3B. OSBP, which harbors a PI4P-binding PH domain and lacks the KKRK polybasic motif, has a lower-confidence LIR and did not interact with GST-LC3B in this assay (Fig. 5D, E). We did not test LC3B binding to ORP8, as it has already been shown to have ATG8-binding capacity and to interact with PIP₂ during lipophagy (Pu et al., 2023). ORP9, a core PITT pathway component, lacks a predicted LIR and, as expected, showed no interaction with GST-LC3B (Fig. S5B, C).

To confirm that the predicted ORP3 sequence constitutes a bona fide LIR, we mutated the WEVV sequence to AEVA (the canonical AXXA substitution; mutations annotated in Fig. 5F). This mutation (ORP3 ΔLIR) completely abolished ORP3 interaction with GST-LC3B after LLOME treatment, without disrupting ORP3 phosphorylation. In contrast, a mutant of the PIP_2_-binding polybasic motif (ORP3 ΔPH) was not phosphorylated and likewise failed to bind GST-LC3B (Fig. 5G–I). Together, these findings suggest that association with membranes is critical for both ORP3 phosphorylation and its interaction with LC3B.

### Structure/function analysis of ORP3 in lysosome repair

To determine whether ORP3 interactions with LC3B and PIP_2_ or its lipid transfer activity are essential for lysosome localization and repair, we used ORP3 mutants ΔLIR, ΔPH, and an ORD domain mutant, (ΔORD) in which two canonical histidines that are critical for PI4P-dependent phospholipid counter-transport are replaced by alanines (de Saint-Jean et al., 2011; Tong et al., 2013) (Fig 5F)

We first assessed whether any domain mutants disrupted ORP3 localization to damaged lysosomes. As expected, WT ORP3 prominently localized to LAMP1/LC3B-positive autolysosomes after 3 hours of LLOME treatment (Fig. 5J, K). Consistent with the GST-LC3B pulldown results, disruption of either N-terminal site; i.e. the LIR or the polybasic region of the PH domain, rendered ORP3 entirely cytosolic, indicating that interactions with both LC3B and PIP₂ are required for recruitment to damaged lysosomal membranes. In contrast, disruption of the PI4P-binding histidine residues of the C-terminal ORD domain did not significantly affect ORP3 recruitment (Fig. 5J, K), indicating that neither PI4P binding nor lipid transfer activity of the ORD domain is required for ORP3 recruitment.

To assess whether lipid transfer activity via the ORD domain is required functionally for lysosome repair, we generated shRNA-resistant versions of the ΔLIR and ΔORD mutants and used these in a Gal3 recovery assay in ORP3-depleted cells. Here we expressed unfused GFP, GFP-WT ORP3 (positive control), GFP-ΔLIR, or GFP-ΔORD at comparable levels (Fig. 5L). As shown in Fig. 5M, N, ORP3-depleted cells expressing only GFP retained numerous Gal3 puncta after 8 hours of recovery. As expected, rescue with WT GFP-ORP3 reduced Gal3 puncta to background levels comparable to shControl cells, and the LIR-deficient mutant, which is not recruited to damaged lysosomes, failed to restore lysosomal integrity. Notably, the lipid transfer-deficient ΔORD mutant likewise failed to restore lysosomal integrity, indicating that removal of PI4P and/or counter-transfer of phosphatidylcholine from the ER to lysophagosomes is critical for recovery from lysosomal damage.

### Lysosome disruption via Cationic Amphiphilic Drugs (CADs) causes ubiquitination, p38 activation, and LC3B–ORP3 interaction

Given that LLOME is a broad lysosome-disrupting agent that can affect lysosomal signaling in multiple ways (Ballabio and Bonifacino, 2020; Radulovic et al., 2026; Nixon, 2024), we sought to distinguish which aspect of lysosomal dysfunction drives ORP3 activation. To this end, we tested compounds that target individual LLOME-induced phenotypes more selectively and assessed their effects on p38 and ORP3 activation (Fig. 6A–B). Inhibitors of mTORC1 (Torin 1), lysosome pH neutralization (NH₄Cl), proteases (E64/PEPA), PI-3 kinase (Wortmannin) or the PI-5 kinase PIKFyve (Vac1) did not induce significant activation of either p38 or ORP3, nor did the TRPML1 agonist (ML-SA1) which drives lysosomal calcium release. In contrast, cationic amphiphilic drugs (CADs) and/or functional inhibitors of acid sphingomyelinase (FIASMAs), such as chloroquine, which cause membrane packing defects through direct interactions with lysosomal membranes or the enzymes that modify membrane lipids (Kaur et al., 2025; Abe et al., 2007; Hinkovska-Galcheva et al., 2021), specifically activated both p38 and ORP3 similarly to LLOME (Fig. 6A-C; Summarized in Fig. S6A, B).

We found that three structurally distinct CADs, the anti-arrhythmic amiodarone (AMD), the tricyclic antidepressant amitriptyline (AMT), and the anti-malarial chloroquine (CHQ), activated p38 and induced ORP3 phosphorylation to varying degrees (Fig. 6A-F). Tetracycline (TTC), which shares structural similarities with CADs but does not cause phospholipidosis or inhibit lysosomal lipases (Abe et al., 2007), activated neither p38 nor ORP3 (Fig. 6D-F). In addition, AMD, AMT, and CHQ each increased cell death in ORP3-depleted cells, while tetracycline did not (Fig. 6G). Importantly, AMD- and AMT-induced ORP3 phosphorylation was blocked by the p38 inhibitor doramapimod (Fig. S6C, D). Consistent with this, p38-dependent ORP3 activation by AMD increased ORP3 binding to GST-LC3B, while the negative control TTC did not (Fig. 6H, I). AMD also induced a corresponding increase in ORP3 to LC3B colocalization (Fig. 6J).

At the doses and timepoints examined, both LLOME and AMD, but not TTC, induced lysosomal ubiquitination, consistent with their capacity to activate p38 and ORP3 (Fig. 6L, M). However, AMD did not cause sufficient membrane disruption to expose lumenal glycans, as indicated by the absence of Gal3 puncta (Fig. 6L, N). Together, these data demonstrate that even modest lysosomal membrane disruption is sufficient to drive lysosomal ubiquitination and ORP3 activation, and to reduce cell survival in the absence of ORP3.

## Discussion

In this study, we identify a critical role for the lipid transfer protein ORP3 in late-stage lysosomal membrane repair. Loss of ORP3, or disruption of its lipid transfer function, leads to defective restoration of lysosome function, increased lipid peroxidation, and reduced cell survival.

Following lysosome disruption by LLOME, ORP3 is recruited to ER–lysophagosome contact sites via a signaling cascade triggered by lysosomal ubiquitination and mediated by TAK1, p38, and, to a lesser extent, IKK. p38-dependent phosphorylation of ORP3 promotes its interaction with the ATG8 protein LC3B, which is required for ORP3 recruitment to damaged lysosomes. We hypothesize that phosphorylation induces a conformational change in the ORP3 N-terminus, exposing a conserved LC3-interacting region (LIR) and PIP₂-binding PH domain to enable recruitment, possibly through a coincidence detection mechanism. We further show that ORP3 phosphorylation and recruitment are triggered by CADs, which induce lysosomal membrane ubiquitination independently of limiting membrane permeabilization, as indicated by the absence of Gal3 recruitment.

We previously demonstrated that ORP3 is recruited to focal adhesions at the plasma membrane in response to Ca^2+^ influx in a PKC-dependent manner, where it establishes ER/plasma membrane contact sites (D’Souza et al., 2020). In that study, we showed that ORP3 extracts PI4P from the plasma membrane in exchange for PC, which we identified as the primary cargo for its lipid-binding ORD domain. Here we show that lysosomal damage triggers ORP3 phosphorylation through a different, ubiquitin/TAK1/p38-dependent mechanism that instead drives its recruitment to lysosomal membranes. Mutants lacking a functional LIR or PH domain failed to be recruited to damaged lysosomes, and while a lipid transfer-deficient mutant was efficiently recruited, it failed to support lysosome repair, strongly suggesting that delivery of ER-derived PC is essential for effective lysosome repair.

### ORP3 in temporal relation to other lysosome associated lipid transfer proteins

As described above, lysosome repair is thought to occur in an ordered series of events (Xun and Tan, Kravic and Meyer reviews). Within minutes of LLOME treatment, lysosomal Ca^2+^ release triggers recruitment of the bridge-like lipid transport proteins (BLTPs) VPS13C (Wang et al., 2025b) and BLTP3A (Hanna et al., 2025 *preprint*) and elements of the ESCRT machinery, which are thought to repair damaged membranes by direct membrane repair and/or micro-lysophagy (Hoyer et al., 2022; Lee et al., 2020). Slightly later (10-40 mins), Ca^2+^-stimulated production of PI4P by PI4K2A initiates recruitment of selected OSBP-related lipid transfer proteins that, like the BLTPs, establish and populate ER/lysosome contact sites. These include ORP9, ORP10 and ORP11, which act together to transfer PS from the ER to lysosomes, as well as OSBP and ORP1L, which transfer cholesterol (Tan and Finkel, 2022; Radulovic et al., 2022). Another BLTP, ATG2, arrives slightly later, by binding to PS (Gao et al., 2010; Tan and Finkel, 2022; Cross et al., 2023). In contrast, ORP3 appears to function at a significantly later stage of lysosome repair (hours vs minutes). Although SPG20-driven K63 ubiquitination and TAK1/p38 activation begins early (5-20 mins after damage induction) (Gahlot et al., 2024; Endo et al., 2025a) (Fig. S3A, B), K63 ubiquitination continues for hours, peaking at ∼3h (Endo et al., 2025a; Papadopoulos et al., 2017). We found that TAK1 inhibition completely blocks both ORP3 phosphorylation and its recruitment to damaged lysosomes without inhibiting LC3B recruitment (Fig. 3A-G)

PC is a phospholipid comprising more than 50% of most eukaryotic membranes (Ali and Szabó, 2023). The bridge-like lipid transfer proteins (BLTPs), such as VPS13C, BLTP3, ATG2 are thought to deliver bulk phospholipids with limited lipid species selectivity, driven primarily by lipid gradients (Hao et al., 2026). Although specific cargo lipids for BLTPs are not well defined, studies of *Drosophila* autophagic membranes lacking ATG2 indicate defects in PE delivery, while PC content of these membranes is not significantly different from wild type (Laczkó-Dobos et al., 2021). PI4P-driven counter-transport by the ORP3 ORD domain may confer directionality and selectivity for PC with specific acyl chain compositions. Notably, monounsaturated fatty acid (MUFA)-containing 18:1 PC was identified as a preferred ORP3 cargo in our previous work (D’Souza et al., 2020). Delivery of MUFA-PC to lysophagosomal membranes may confer protection against lipid peroxidation following lysosomal membrane permeabilization (LMP), as MUFA-containing phospholipids are less susceptible to oxidative damage than polyunsaturated species (Cortie and Else, 2015; Qiu et al., 2024; Firsov et al., 2025; Park et al., 2025). Consistent with this possibility, acute amiodarone CAD treatment in rats led to a significant hepatic enrichment of 18:1 PC (Thu et al., 2025 *preprint*), and PI4KIIα-driven ORP6 recruitment in LLOME-treated neuronal HT22 cells specifically increased MUFA 18:1 PC on lysosomal membranes (Zhang et al., 2026). Finally, the localization of ORP3-positive lysophagosomes proximal to the nucleus may be associated with PC synthesis machinery, as the rate limiting enzyme of PC synthesis, CTP:phosphocholine cytidylyltransferase α (CCTα), localizes to nuclear/ER continuous membranes, which are rich in 18:1 PC (Hunt et al., 2001; Cornell and Antonny, 2018; Haider et al., 2018). Studies in yeast have determined that de novo PC generated by CCTα is crucial for late stages of autophagy completion (Polyansky et al., 2022)

### Termination of ORP3 activity by deubiquitination of lysosomal membranes

Our data suggest that recruitment of the VCP/p97 de-ubiquitinating complex at late stages of lysosome repair mediates termination of the TAK1/p38-dependent activation of ORP3 via clearance of K63 ubiquitin. We hypothesize that the level of K63 ubiquitin on lysosomal membranes correlates with the extent of repair, diminishing over time as repair proceeds, leading to a parallel decrease in TAK1/p38 signaling. This equates to a TAK1 “on” switch, and a VCP/p97 “off” switch for ORP3 phosphorylation. Reinders et al recently reported that deubiquitination is mediated by a complex of VCP/p97 and the deubiquitinase Ataxin-3 (Reinders et al., 2025). Interestingly, Ataxin-3, like ORP3, is recruited to PIP_2_ positive lysosomes that have not been acidified within 3h of LLOME treatment, suggesting that they are still undergoing repair. We propose that these non-acidified, ORP3/LC3B/LAMP1-positive structures represent autophagosome-engulfed lysosomes that have not been resolved by ESCRT-or PITT-mediated repair and have not yet undergone lysophagy (Reinders et al., 2025), and may be involved in Autophagic Lysosome Regeneration (ALR).

Taken together, our data support a model in which lysosomal damage triggers ubiquitin-dependent phosphorylation of ORP3, which is required for its recruitment through the combined activities of its N-terminal LIR and PH domains. In this model, PIP₂ production, which occurs late in the repair process, is essential for complementing the LIR-LC3B interaction. This distinguishes ORP3 recruitment from PI4P dependent PITT recruitment in both time and space. We hypothesize that ORP3-mediated transfer of PC from the ER to the lysosome associated autophagic membranes is required for either completion of the repair process or progression through lysophagy.

## Methods

### Cell Culture

HeLa, HEK-293T, and HEK293FT cells were obtained from the ATCC. All cells were cultured in 10% FBS, 1% Pen/Strep Dulbecco’s Modified Eagle’s Medium (DMEM) +High Glucose, +NaPyruvate, +Glutamine). Cells were grown in standard conditions, 37°C and 5% CO_2_. All media were carefully equilibrated in an incubator before changes. Cells were routinely checked for mycoplasma contamination.

Transfection of HEK293T and HEK293FT cells was performed using Polyjet (Signagen) according to the manufacturer’s instructions. For imaging, HeLa cells were transfected with Fugene4k (Promega) according to the manufacturer’s instructions.

### Plasmids

All constructs are based on human sequences. GFP-ORP3 and GFP-ORP3 ΔPH (60-KKRK-63 to 60-IAAI-63) and shRNA-resistant GFP-ORP3 constructs were prepared previously (D’Souza), using plasmids originally provided by Vesa Olkonnen (University of Helsinki, Finland). GFP-ORP11, GFP-ORP6, and GFP-ORP9L were gifts from Li Yang and Hongyuan (Robert) Yang (University of New South Wales, Sydney, Australia). pEGFP-C1-OSBP was a gift from David Castle (University of Virginia). pEGFP-C1-PLCδ (Addgene #21179), Halo-Sec61-C-18 (Addgene #123285), CMV-LAMP1-HaloTag (Addgene #164209), pHTN-HaloTag_RAB7A:1-207 (Addgene #211095) were from Addgene. All LC3B constructs were also from Addgene: RFP-LC3B (Addgene #200943), pEGFP-LC3 (Addgene #24920), GST-LC3B- pGEX4T1-GST-LC3B (Addgene # 216836).

### Mutagenesis

All ORP3 mutants (ΔHH and ΔLIR) were generated from both WT GFP-ORP3 and GFP-ORP3 shRNA-resistant constructs described above using a Q5 site directed mutagenesis kit (Promega) using NEB designed primers. ΔLIR (33-WEVV-36 to 33-AEVA-36) mutants were generated using oligos FWD 5’ GAAGTGGCGGAAGGACTGAGGGGG 3’ and REV 5’ CGCGCTGTCCTGTCGACTTCCTTGC and ΔHH (630-HH-631 to 630-AA-631) mutants were generated using oligos FWD 5’ ACAGGTCAGCgcggcgCCGCCTATCTCTG 3’ and REV 5’ TCTGAAAAAAACTGGAAGC 3’.

### Antibodies

The following primary antibodies were used in this study: mouse anti-ALIX (Santa Cruz Biotechnology sc-25279, RRID:AB_627656), mouse anti-Galectin3 (Santa Cruz Biotechnology sc-32790, RRID:AB_627657); mouse anti-LAMP1 (DSHB Cat# H4A3, RRID:AB_2296838); rabbit anti-LAMP1 (Cell Signaling Technology Cat# 9091, RRID:AB_2687579); mouse anti-ORP3 (Santa Cruz Biotechnology Cat# sc-398326, RRID:AB_3699205); rabbit anti-IкB (Cell Signaling Technology Cat# 9242, RRID:AB_331623); rabbit anti-p38 MAPK (Cell Signaling Technology Cat# 9212, RRID:AB_330713); rabbit anti-phospho-p38 MAPK (Thr180/Tyr182) (Cell Signaling Technology Cat# 9215, RRID:AB_331762); mouse anti-Actin (Millipore Cat# MAB1501, RRID:AB_2223041); rabbit anti-beta-Actin (Proteintech Cat# 81115-1-RR, RRID:AB_2923704); rabbit anti-mCherry (Thermo Fisher Scientific Cat# PA5-34974, RRID:AB_2552323); rabbit anti-GFP (Proteintech Cat# 50430-2-AP, RRID:AB_11042881); mouse anti-GFP (Proteintech Cat# 66002-1-Ig, RRID:AB_11182611); mouse anti-FK2 (Enzo Life Sciences Cat# BML-PW8810-0500, RRID:AB_2051891)

### shRNA

Lentiviral shRNA vectors (pLKO.1) targeting ORP3-TRCN0000157183 (GCCTTTGCCATATCAGCGTAT) were gifts from Johnny Ngsee (University of Ottawa, Canada) and used as described previously (D’Souza 2020). Empty pLKO.1 vector (Addgene) was used as control for all knockdown experiments. Lentivirus was produced in HEK293FT cells as previously described (D’Souza 2020). HeLa cells were transduced with virus for 24h, selected using 2.5ug/ml of puromycin, and analyzed 3-4 days later

### Chemicals, inhibitors, and stimuli

Inhibitors: Target-Drug Dose (Source): Ulk1/2i- MRT-68921, 1μM (Cayman Chem #19905); TBK1i- MRT67307, 1μM (Cayman Chem #19916); IкK1/2(α/β)i- TPCA-1, 10μM (Cayman Chem #15115); TAK1i- (5Z)-7-Oxozeaenol, 20μM (Cayman Chem #17459); p38i-Doramapimod, 200nM (Cayman Chem #10460); All p38 isoforms. *Used for all p38i conditions unless otherwise noted; p38i- SB202190, 1μM (Cayman Chem #10010399) p38 α/β isoforms; p38i- VX-702, 10μM (Cayman Chem #13108); p38 α/β isoforms; AmpKi- Dorsomorphin (HCl), 10μM (Cayman Chem #21207); ISRIBi (Integrated Stress Response)- Trans-ISRIBi, 200nM (Cayman Chem #16258); CK2i (Casein Kinase 2)- IQA, 20μM (Cayman Chem #35223); LRRK2i- LRRK2-IN-1, 1μM (Cayman Chem #18094); PKCi-Go 6983, 10μM (Tocris #2285); ATMi- Ku-55933, 2μM (Cayman Chem #16336); DNA-PKi- NU 7441, 2μM (Cayman Chem #14881); PI3Ki- wortmanin, 1μM (Cayman Chem 10010591); TRPML1i-ML-SI1, 10μM Gift from Bettina Winckler (University of Virginia); JNKi- SP600125, 10μM (Cayman Chem #10010466); ERK1/2i- GDC-0994, 10μM (Cayman Chem #21107); VCP/p97i-NMS-873, 1μM (Cayman Chem #17674).

Stimuli: LLOME- L-leucyl-L-leucine methyl ester hydrobromide, 0.5mM (unless otherwise noted) (Sigma #L7393) LOME- L-leucine methyl ester hydrobromide, 2mM (Sigma #L1002); Thapsigargin (SOCE Store operated Calcium Entry and PKC activation), 1μM (Tocris #1138); Torin1 (MTOR inhibitor and macro-autophagy inducer), 500nM Gift from Winckler Lab; Vacuolin1 (PIKfyve inhibitor and PITT inducer), 1μM (Cayman Chem #20425); E64 (protease inhibitor), 50μM (Cayman Chem #10007963) Pepstatin-A (protease inhibitor), 10μg/ml (Cayman Chem ##9000469). N-Acetyl Cysteine (Antioxidant), 2mM (Sigma #A9165); MG-132 (proteasome inhibitor), 10μm (Adooq Bioscience #A11043); NH_4_CL_2_ (lysosomal pH neutralization), 20mM (Sigma # 254134) CADs: each; Amiodarone(HCl) 50μM for 3h for biochemistry and imaging, 20μM for the overnight survival assay (Cayman Chem #15213); Amitriptyline(HCl) 3h 50μM for biochemistry, 20μM for the survival assay (Cayman Chem #15881); Chloroquine (2H_3_PO_4_) 100μM for 3h biochemistry, 20μM for the survival assay; Tetracycline (HCl) 50μM for 3h for biochemistry and imaging, 20μM for the survival assay (Cayman Chem #14328)

### Immunofluorescence microscopy

HeLa cells grown on fibronectin-coated glass coverslips were quickly rinsed in room temperature phosphate buffered saline (PBS) and fixed in pre-warmed (37°C) 4% paraformaldehyde in PBS on the benchtop for 12 minutes. Cells were washed 3x with PBS before subjecting to antibody staining.

Cells were permeabilized in 0.2% Triton-X 100 in PBS for 10 minutes and blocked in 5% BSA in PBS for 30 minutes. Primary antibodies were diluted in 1% BSA in PBS and incubated for 1h at room temperature. Cells were washed 3 times in PBS, before being stained with secondary antibodies in 1% BSA in PBS, again for 1h at room temperature. Secondary antibodies were chosen based on species compatibility and fluorophore (488, 568, 647nm). Cells were washed 3 times in PBS, before mounting with Prolong Gold mounting media.

### Cell survival assay

HeLa cells were plated on 96-well, black bottom, Cell-gro plates (Corning) overnight. LLOME or indicated CADs were added the next day, overnight. Alternatively, LLOME was added for 1h before being washed out and chased in fresh media, as indicated. After18-20 hours, cells were rinsed with PBS, and fixed in pre-warmed 37°C 4% paraformaldehyde/PBS at room temperature for 12 minutes. Fixed cells were washed 3x with PBS before staining with Alexa488-Phalloidin (SCBT #sc-363791) in 0.2% Triton-X, 1% BSA/PBS for 1hr. Cells were washed 3x in PBS and Alexa488 fluorescence was measured using a Cytation 1 fluorescence microplate reader (Biotek).

### Live Imaging

Cells were grown on fibronectin-coated 35mm glass bottom dishes (Mattek). Cells were transfected and subjected to imaging the next day in complete phenol-red free DMEM (Gibco) imaging media supplemented with 10mM HEPES, pH 7.4. Cells were allowed to equilibrate in a stage-top incubator (Tokai) for 10 minutes before imaging. To label lysosomes, 40ug/ml 70kD Alexa 633-dextran (Biotium 80141) was fed overnight, washed 3x with fresh media and chased for 5h for terminal compartment labeling before cell treatments and imaging. Lysotracker-Red (Molecular Probes L-7528) or Lysoview633 (Biotium #70058-1) were added to cells 40 minutes prior to imaging. To detect lipid peroxidation, NBD-Pen (TargetMol #1955505-54-0)(0.2uM) was added simultaneously with LLOME or for the same amount of time (5h) for untreated cells (DMSO). Cells were washed 1x with pre-warmed media, and replaced with pre-warmed imaging media for microscopy. HaloTag-labeled organelle markers (Halo-LAMP1, Halo-Rab7, and Halo-Sec61) were visualized with HaloTag ligand JF669- provided by the Lavis lab at Janelia Farm (Ashburn, Virginia, USA). HaloTag dye was added to cells 30 min before imaging, no media change required.

### Microscopy

Cells were imaged using a Nikon AX-R laser scanning confocal microscope (Nikon Instruments) fitted with a 100x Oil 1.4 NA PlanApo TIRF DIC N2 objective. For super-resolution imaging, a Nikon NSPARC module was used. Laser power and gain settings were identical within experiments for all conditions. For live imaging, a Tokai Hit stage top incubator was used to maintain cells at 37°C.

For spinning disc microscopy, we used a Nikon Ti2S motorized microscope equipped with a Yokogawa CSU-X1 spinning disc and 60x Oil PLAN APO λD 1.42 NA, DIC objective. For live super-resolution imaging we used a Nikon W1-SoRa spinning disc confocal microscope fitted with a CFI60 60X Oil PLAN APO λD, 1.42 NA, DIC objective.

### Cell lysis and immunoprecipitation

Cells were rinsed in ice cold PBS and lysed in ice cold immunoprecipitation (IP) buffer (HeLa cells: 0.5% NP-40, 20mM Tris pH 7.4, 100mM NaCl; for HEK293T cells:1% Triton-X 100, 20mM Tris pH 7.4, 100mM NaCl) supplemented with 1:100 HALT protease/phosphatase inhibitor cocktail (Thermo fisher scientific), directly on plates. Cell Lysates were scraped into 1.5ml Eppendorf tubes and rotated for 20 minutes at 4°C before centrifugation at 12,000 x g for 10minutes at 4°C. Aliquots of post-nuclear supernatants were diluted with 5x Laemmli SDS-PAGE buffer, boiled 5 minutes, and run on 4-20% gradient gels prior to processing for western blots.

For immunoprecipitation, 500ul of post-nuclear supernatants were precleared with equivalent amounts of protein A/G agarose (Gene DEPOT #P9203-015) for 30 minutes. Cleared lysates were mixed with 1ug anti-GFP antibody for 1h, followed 15ul protein A/G agarose 50% slurry for 1h. Immunoprecipitates were eluted in 15ul 2x Laemmli buffer and associated input lysates were resolved on SDS-PAGE gels prior to immunoblotting.

### Pulldowns

For GST pulldowns, 4ug of unfused GST or GST-LC3B in 15ul glutathione-agarose slurry (Pierce #16100) was added to each lysate for incubation overnight. Agarose pellets were washed with IP buffer 3 times, eluted in 15ul 2x Laemmli buffer, and subjected to SDS-PAGE for Ponceau staining and immunoblotting.

### Phosphatase Assay

For the Lambda Phosphatase dephosphorylation assay (SCBT #sc-200312A), HeLa cells were treated with either LLOME or thapsigargin for 1h and lysates were subjected to enzymatic de-phosphorylation. Briefly, after lysis and a post-nuclear spin, a supernatant sample was removed to keep on ice as a negative control. To the remainder, 10X phosphatase buffer and 2mM MnCl_2_ was added. This was split, with one half given phosphatase inhibitor cocktail (mock control), the other half given lambda phosphatase enzyme, and each incubated at 30°C for 30 min. Samples were then subjected to SDS-PAGE followed by western blotting.

### Immunoblotting

Protein samples were resolved on 4–20% SDS-gradient gels (BioRad, Hercules, CA), then transferred to nitrocellulose membrane (Millipore, Burlington, MA). Membranes were blocked with Intercept blocking buffer (Li-Cor), Tris-buffered saline (TBS)-based when probing phospho-epitopes, or PBS-based for all others. Primary and secondary antibodies were incubated for 1h at room temperature in blocking buffer with 0.05% Tween-20. Detection was carried out using a LI-COR Odyssey infrared scanning system using fluorescently labelled secondary antibodies. Quantification was performed with Image Studio Lite using the densitometry tool. The value of the precipitates relative to the input was normalized for plotting. p38 activity was measured as the ratio of phospho-p38 to total p38 and normalized to untreated or LLOME-treated cells as indicated.

For ORP3 and LC3B band shift ratios, individual peaks were manually measured using ImageJ on band profiles generated using Image Studio Lite. The ratio of the upper to lower band (ORP3), or the lower to upper band (LC3B) were measured and normalized to untreated cells.

IкB levels were measured via densitometry relative to an actin loading control.

### Recombinant Protein Purification

GST or GST-LC3B were expressed in *Escherichia coli* BL21. Bacteria were grown at 37°C to an OD600 of 0.6 to 0.8, before expression was induced with 0.5mM isopropyl β-D-thiogalactopyranoside (IPTG) for 2-4 h at 30C. Cells were harvested via centrifugation at 6000g for 20 min. Pellets were washed with ice-cold PBS before resuspension in Buffer A (25mM Tris HCl pH 7.4, 1M NaCl, 0.5mM EDTA, 1mM DTT, 0.1% TX-100, Complete protease inhibitors (Roche) and phenylmethylsulfonyl fluoride (PMSF, 1 mM)). Cells were lysed by sonication, and cleared via centrifugation at 10,000 x g for 10min at 4°C. Pre-equilibrated glutathione-sepharose beads were added to supernatants and incubated with rocking for 30-45 minutes at room temperature. Bead pellets were washed 3x with ice-cold PBS, and resuspended in Buffer A to a 50% slurry. Protein concentration of the bead slurry was quantified by Coomassie Blue staining of SDS-PAGE gels using linear BSA reference standards.

### Image analysis

Images were background subtracted uniformly within experiments. Colocalization analysis was performed using NIS Elements AR software, using the built-in colocalization tool to determine Pearson’s or Mander’s colocalization coefficients, and mean intensity using the ROI measurement tool on the indicated channels.

To quantify Gal3-, GFP-ORP3- or, GFP-ORP11- positive lysosomes, LAMP1 positive ROIs were generated using automated SPOT detection (NIS elements AR) and assessed for the presence of labeled protein above background threshold in the ROIs.

Lysosomal lipid peroxidation was quantified by assigning ROIs to Lysotracker-positive lysosomes using automated SPOT detection (NIS elements AR) and quantifying NBD-Pen intensity above assigned background threshold in the ROIs.

## Statistical Analysis

Graphpad Prism 11.0.1 and Excel were used for all statistical analysis. Shapiro-Wilk normality test was used to differentiate between normal/non-normal distributions, and appropriate statistical tests were chosen based on the results of the normality test and number of comparisons. Mann Whitney U test was used to test for significance between two non-parametric groups. In the case of multiple comparisons, the Kruskal-Wallis one-way analysis of variance was used along with Dunn’s multiple comparison tests, or Fieldman’s test for matched observations. For normal data, student’s ‘t’ test was used for single comparisons, or One-way ANOVA for multiple comparisons. SD and mean are shown unless otherwise indicated.

Significance indicated as ns. > 0.05, * <0.05, **<0.01, ***<0.0005, and ****<0.0001

## Supplemental Material

Supplemental data includes relevant data corresponding to the indicated figure. These include additional evidence and avenues of the mechanism of ORP3 activation, comparisons of super-resolution to conventional confocal microscopy techniques, and further comparison of human ORP protein sequences in relation to n-terminal LC3 and PIP2 binding.

## Data Availability Statement

The data are available from the corresponding author upon request.

## Acknowledgements

The authors thank Michael Junyup Lim, Sophia Link, Lindsey Haag, Sophia Spinu, and Danny Cao for technical assistance. We thank Li (Emma) Yang, Hongyuan (Robert) Yang, and Vesa Olkonnen for ORP constructs and David Castle (University of Virginia) for OSBP and PLCδ-PH plasmids. We also thank Laura Newman (University of Virginia) for use of her spinning disc confocal microscope and Elias Spiliotis (University of Virginia) for use of the SoRa super-resolution spinning disc microscope. Finally, we thank David Castle, Laura Newman and Bettina Winckler for thoughtful discussions and for reading the manuscript. Cartoons were created with BioRender (Biorender.com)

---

**Figure S1.**
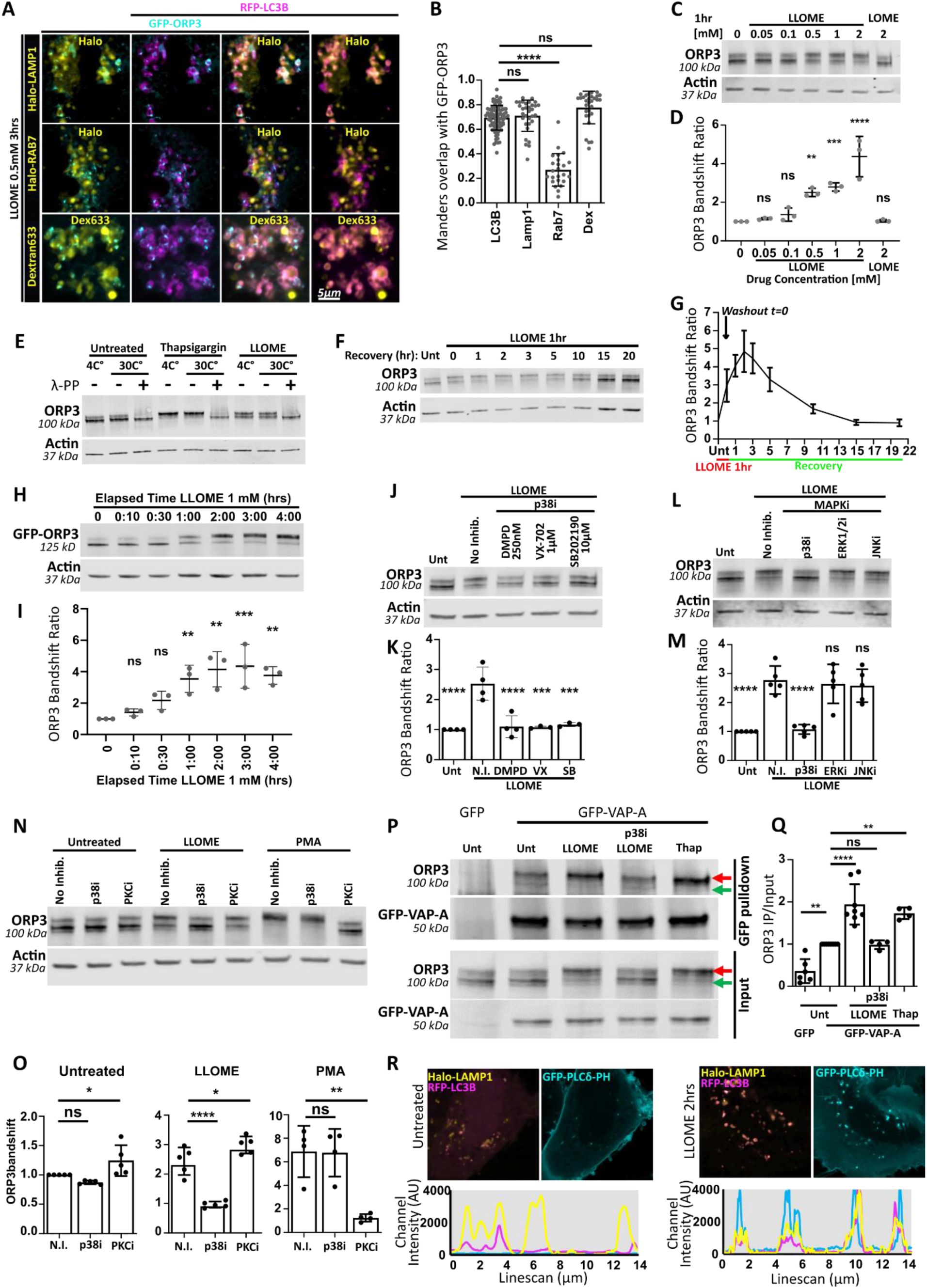
**(A)** Live imaging of cells co-expressing GFP-ORP3 and RFP-LC3B with Halo-LAMP1 (top row), Halo-Rab7 (middle row) or incubated with far red Alexa633-dextran after 3h treatment with LLOME. **(B)** Quantification of Manders overlap of each probe with GFP-ORP3 in **A**. 20-30 cells per condition over 1 experiment. Paired mixed effects analysis with Dunnet’s correction compared to LC3B. **(C)** HeLa cells were treated with increasing doses of LLOME (or LOME, last lane) for 1h, lysed, and immunoblotted for endogenous ORP3. (**D**) Quantification of band shift ratio in **A**. n=3 experiments. One-way ANOVA with Dunnett’s correction tests vs untreated/vehicle (dose 0 mM). **(E)** HeLa cells were untreated or treated with thapsigargin or LLOME for 3h. Lysates were then untreated or incubated with lambda phosphatase at either 30°C, or 4°C (negative control). Collapse of the ORP3 doublet in the presence of phosphatase is indicative of phosphorylation. (**F**) Recovery of ORP3 LLOME-induced band shift after LLOME treatment for 1h followed by washout. (**G**) Quantification of 3 to 5 experiments represented in **F**. (**H**) GFP-ORP3 expressed in HEK293T cells yields a similar time course of activation/deactivation as endogenous ORP3 in HeLa cells wen treated with 1mM LLOME. (**I)** Quantification of **H**. n=3 One-way ANOVA relative to untreated (time 0). **(J)** HeLa cells were treated with LLOME for 3 h in the absence or presence of 3 different p38 MAPK inhibitors, then immunoblotted for endogenous ORP3. **(K)** Quantification of ORP3 band shift in **J**. n=3 to 4 One-way ANOVA relative to LLOME without inhibition. **(L)** Cells were incubated with inhibitors of p38 (doramapimod, p38i), ERK1/2 (GDC-0994) or JNK (SP600125) and LLOME, then immunoblotted for endogenous ORP3. **(M)** Quantification of **L**. n=5 One-way ANOVA relative to LLOME without inhibition. **(N)** Comparison of inhibition of ORP3 band shift with LLOME vs PMA in the presence of p38 or PKC inhibitors. **(O)** Quantification of **N**. n=4-5 experiments per condition, One-way ANOVA relative to no inhibitor (N.I.) **(P)** HeLa cells transiently expressing GFP-VAPA were treated with the indicated inhibitors/stimuli for 2h, subjected to anti-GFP immunoprecipitation and assessed for co-immunoprecipitation of endogenous ORP3 with GFP-VAPA by immunoblotting. Green arrowhead indicates lower, inactive ORP3 band, and the red arrowhead indicates active, phosphorylated ORP3. **(Q)** Quantification of **P**. n= 4 to 7 independent experiments per condition. One-way ANOVA with Dunnett’s correction tests vs untreated. (**R**) Cells co-expressing the PIP2 biosensor GFP-PLCδ-PH, RFP-LC3 and Halo-LAMP1 were untreated (left) or treated with LLOME for 2h (right). Representative images and line scans of the indicated constructs are shown.

**Figure S2.**
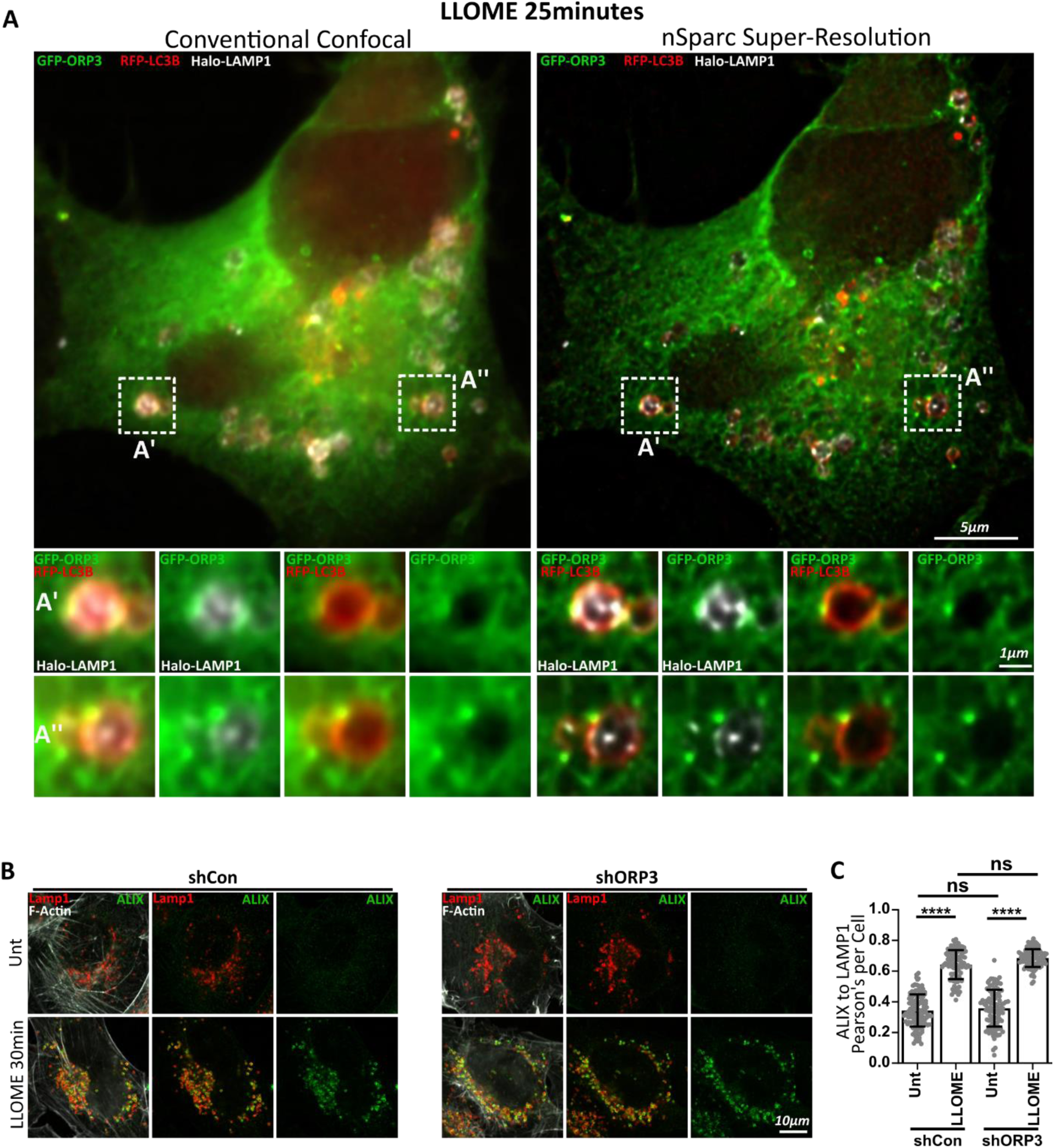
**(A)** Comparison of standard confocal imaging compared to the NSPARC super-resolution mode of the same microscope and cell. Insets for A’ and A” are boxed, showing open-ended ORP3- and LC3B-positive lysophagosomes. **(B)** shCon vs shORP3 HeLa cells were treated with LLOME for 30 minutes and immunostained for the early lysosome damage marker ALIX and LAMP1. **(C)** Quantification of **B**. n=100-120 cells over 2 experiments. Kruskal Wallace test with Dunn’s correction

**Figure S3.**
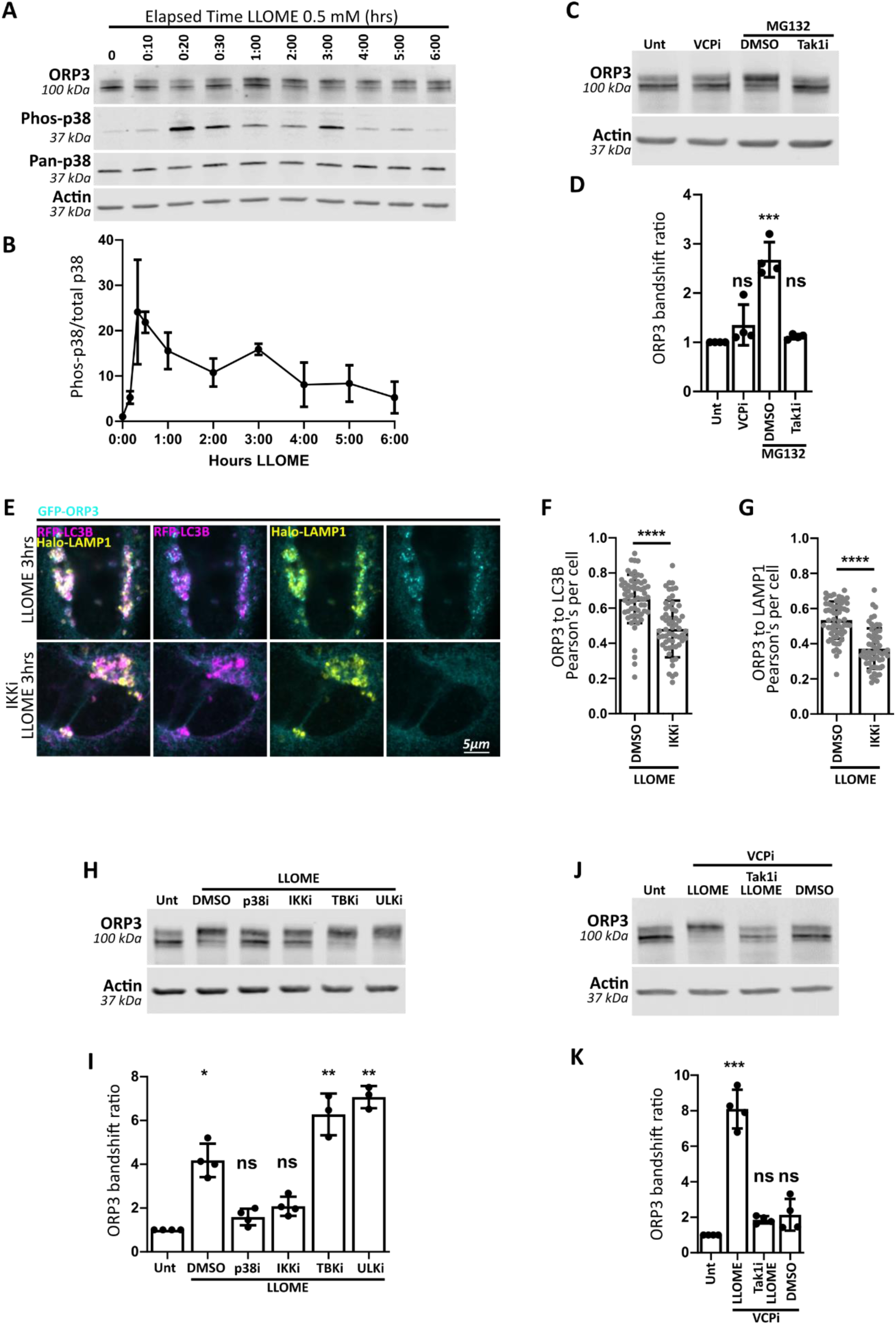
**(A)** Immunoblot of HeLa cells treated with LLOME for the indicated times, probed for endogenous ORP3. The same blot was probed for phospho-p38 and total p38, with actin as a loading control. **(B)** Quantification of phospho-p38 relative to total over the LLOME time course in **A**. Note the secondary peak (∼3hrs) occurring after the initial peak (<30 min). **(C)** MG132, which causes accumulation of K63 ubiquitin, activates ORP3 in a TAK1-dependent manner. **(D)** Quantification of ORP3 band shift in **C**. n=4 experiments, Kruskal Wallace Test. **(E)** HeLa cells transiently expressing GFP-ORP3 and RFP-LC3B were treated with LLOME for 3 h, in the absence (top) or presence (bottom) of the IKK inhibitor *TPCA-1* (IKKi) 30 min prior to LLOME treatment. Cells were then fixed and immunostained for LAMP1. **(F, G)** Quantification of colocalization of ORP3 with LC3B or LAMP1, respectively. N=∼60 cells per condition over 3 experiments. Unpaired t-test. **(H)** HeLa cells were treated with LLOME for 3 h, and/or treated with inhibitors of additional kinases downstream of TAK1. p38 inhibitor doramapimod (p38i), IKK inhibitor TPCA-1 (IKKi), TBK1 inhibitor MRT67307 (TBK1i), and ULK1/2 inhibitor MRT-68921 (ULKi). Endogenous ORP3 band shift was inhibited by p38i, partially inhibited by IKKi, and exacerbated by TBK1i and ULKi. **(I)** Quantification of ORP3 band shift in **H**. n=3-4 experiments, Kruskal Wallace Test. **(J)** ORP3 immunoblot of HeLa cells treated as indicated for 5 h. The VCP inhibitor had little effect in the absence of lysosomal damage induced by LLOME. **(K)** Quantification of endogenous ORP3 activation in **I**. n=4 experiments, Kruskal Wallace Test.

**Figure S5.**
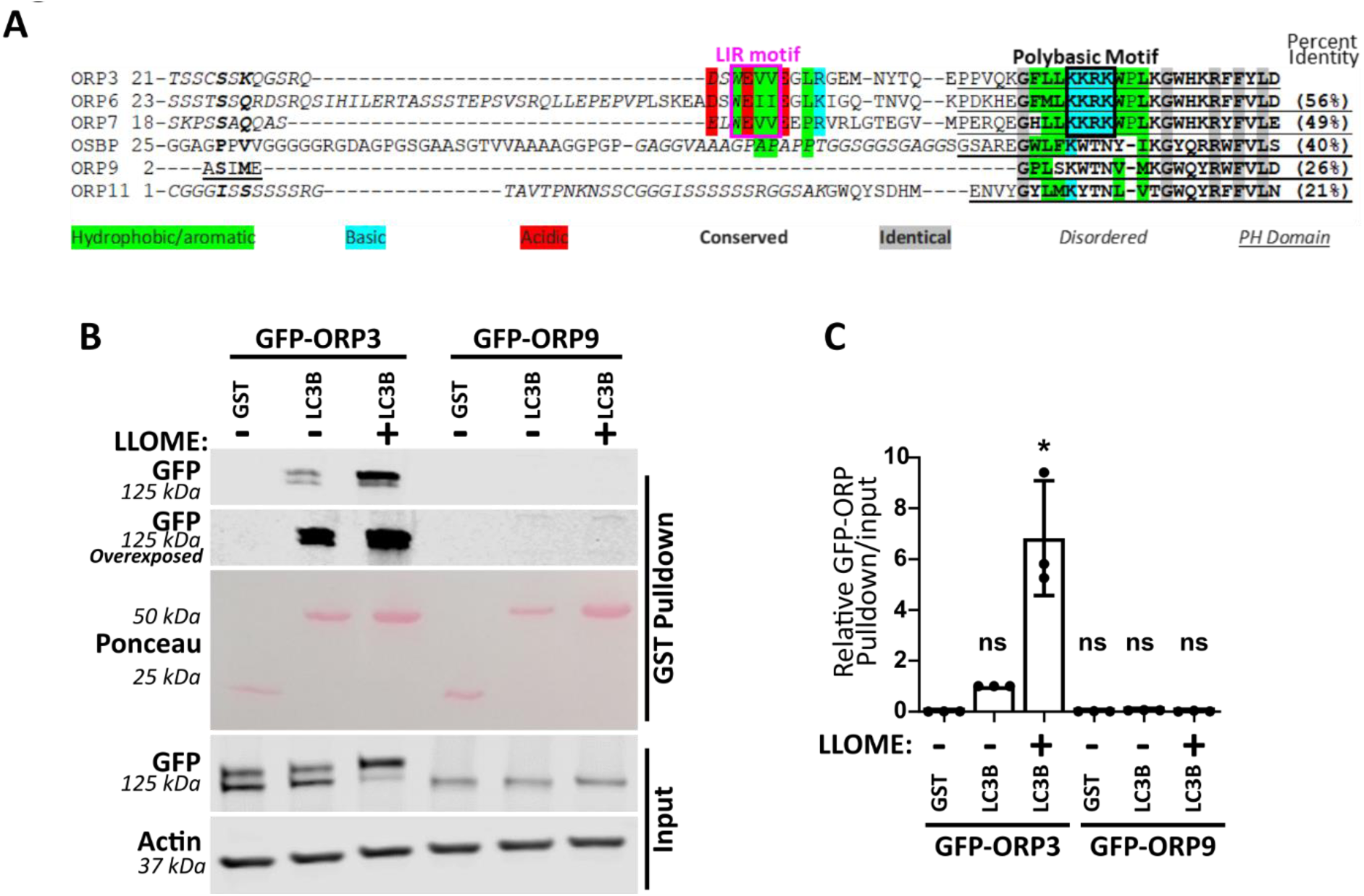
**(A)** Constraint-based Multiple Alignment of the conserved LIR and polybasic motif in aligned human lysosome-associated ORP proteins. OSBP subfamily III: ORP3, ORP6, ORP7; OSBP subfamily I: OSBP; OSBP subfamily V: ORP9; OSBP subfamily VI: ORP11. Conserved features are labeled. **(B)** HEK293T cells transiently expressing GFP-ORP3 or GFP-ORP9, were left untreated or treated with LLOME, lysed and incubated with either GST alone or GST-LC3B. Precipitates were immunoblotted for GFP (**C**) Quantification of the pulldown shown in **B**. N=3 experiments. Friedman Test.

**Figure S6.**
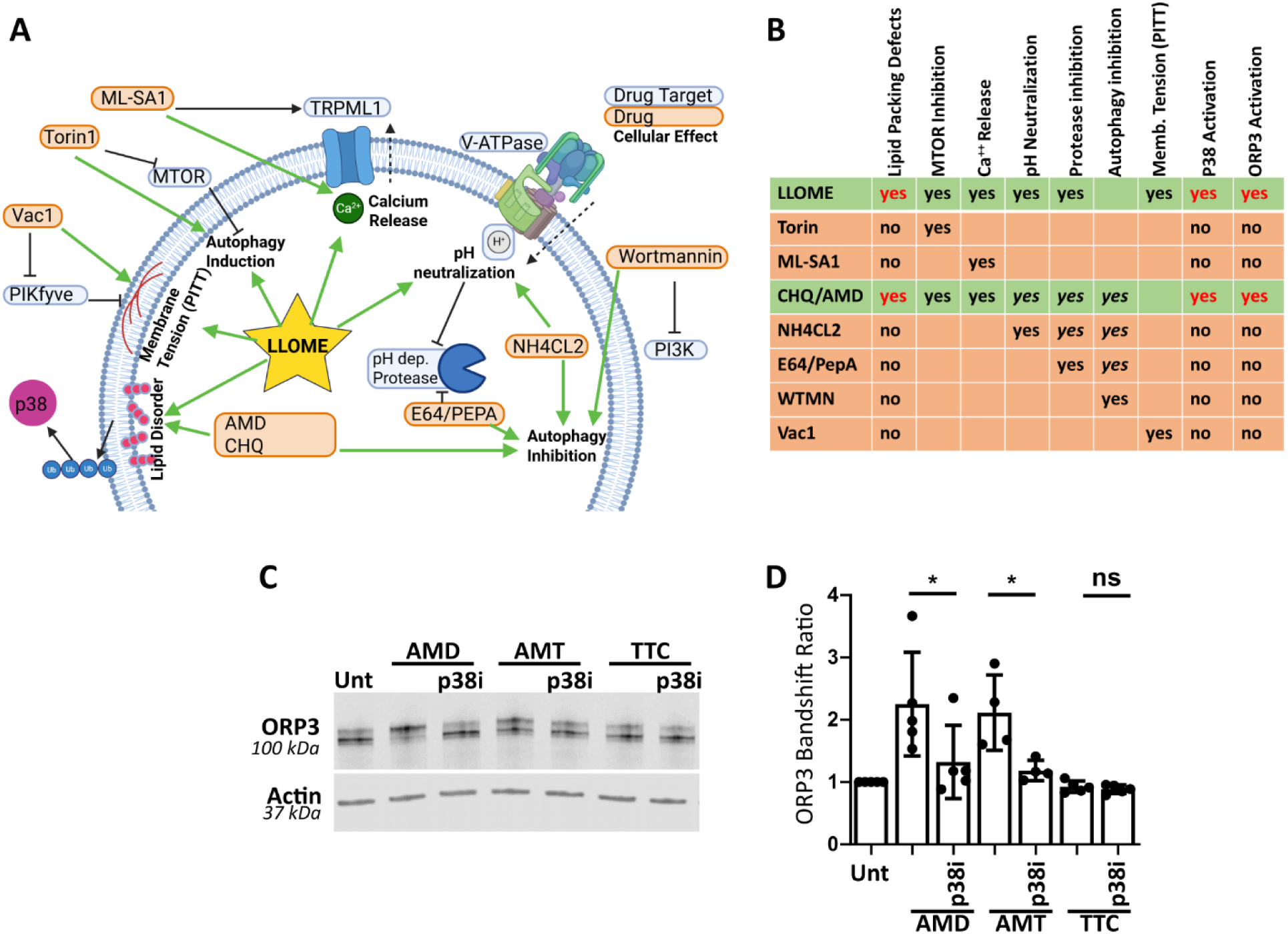
**(A)** Model of how different perturbations affect lysosome function. Created with BioRender. **(B)** Table summary of **A**. **(C)** HeLa cells were treated with the indicated CAD for 3h, in the presence or absence of doramapimod (p38i) and immunoblotted for endogenous ORP3. **(D)** Quantification of ORP3 band shift ratio in **J**. n=4 to 5 experiments, One-way ANOVA with Sidak’s correction.

